# The effect of *Dnaaf5* gene dosage on primary ciliary dyskinesia phenotypes

**DOI:** 10.1101/2023.01.13.523966

**Authors:** Amjad Horani, Deepesh Kumar Gupta, Jian Xu, Huihui Xu, Lis del Carmen Puga-Molina, Celia M. Santi, Sruthi Ramagiri, Steven K. Brennen, Jiehong Pan, Tao Huang, Rachael M. Hyland, Sean P. Gunsten, Shin-Cheng Tzeng, Jennifer M. Strahle, Pleasantine Mill, Moe R. Mahjoub, Susan K. Dutcher, Steven L. Brody

**Affiliations:** Department of Pediatrics, Washington University School of Medicine, St. Louis, Missouri, USA; Department of Cell Biology and Physiology, Washington University School of Medicine, St. Louis, Missouri, USA; Department of Medicine, Washington University School of Medicine, St. Louis, Missouri, USA; Department of Obstetrics and Gynecology, Washington University School of Medicine, St. Louis, Missouri, USA; Department of Neurosurgery, Washington University School of Medicine, St. Louis, Missouri, USA; Donald Danforth Plant Science Center, St. Louis, MO, USA; MRC Human Genetics Unit, University of Edinburgh, Edinburgh, UK; Department of Genetics, Washington University School of Medicine, St. Louis, Missouri, USA

**Author notes:** **Co-corresponding authors:** Corresponding authors: Amjad Horani, MD, Division of Pediatric Allergy, Immunology, and Pulmonary Medicine, Department of Pediatrics, 660 South Euclid Avenue, Mailbox 8116, St. Louis, Missouri, 63110., Facsimile, Steven Brody, MD, Division of Pulmonary and Critical Care, Department of Medicine, 660 South Euclid Avenue, Mailbox 8052, St. Louis, Missouri, 63110., Facsimile: 314362-8980.

**Keywords:** Ciliopathy, cilia, dynein, primary ciliary dyskinesia

## Abstract

DNAAF5 is a dynein motor assembly factor associated with the autosomal heterogenic recessive condition of motile cilia, primary ciliary dyskinesia (PCD). The effects of allele heterozygosity on motile cilia function are unknown. We used CRISPR-Cas9 genome editing in mice to recreate a human missense variant identified in patients with mild PCD and a second, frameshift null deletion in *Dnaaf5*. Litters with *Dnaaf5* heteroallelic variants showed distinct missense and null gene dosage effects. Homozygosity for the null *Dnaaf5* alleles was embryonic lethal. Compound heterozygous animals with the missense and null alleles showed severe disease manifesting as hydrocephalus and early lethality. However, animals homozygous for the missense mutation had improved survival, with partial preserved cilia function and motor assembly observed by ultrastructure analysis. Notably, the same variant alleles exhibited divergent cilia function across different multiciliated tissues. Proteomic analysis of isolated airway cilia from mutant mice revealed reduction in some axonemal regulatory and structural proteins not previously reported in *DNAAF5* variants. While transcriptional analysis of mouse and human mutant cells showed increased expression of genes coding for axonemal proteins. Together, these findings suggest allele-specific and tissue-specific molecular requirements for cilia motor assembly that may affect disease phenotypes and clinical trajectory in motile ciliopathies.

**Brief Summary:** A mouse model of human DNAAF5 primary ciliary dyskinesia variants reveals gene dosage effects of mutant alleles and tissue-specific molecular requirements for cilia motor assembly.

## Introduction

Motile cilia are highly specialized organelles that transport fluids along the surface of the airway, brain ventricle, and fallopian tube, and propel sperm. Over two thousand genes function in the assembly and structure of motile cilia (1). It is not surprising that pathologic variants in over 50 genes are known to be causative of the human motile ciliopathy called primary ciliary dyskinesia (PCD) (2). Classic features of PCD directly reflect dysfunction of organs bearing motile cilia, including chronic upper and lower respiratory tract infection, bronchiectasis, and reduced fertility. Laterality defects may be present and associated with cardiac malformations, owing to the function of motile cilia present in the embryonic node during early development. Despite essential cellular functions of motile cilia, clinical features of patients with PCD vary. Mild disease without the classic features of PCD is increasingly recognized in a subset of patients (3, 4).

Limited epidemiology suggests broad genotype-phenotype relationships for several common PCD genes. For yet undetermined reasons, severe disease is associated with variants in some genes (e.g., *CCDC39, CCDC40*) and mild disease in others (e.g., *DNAH9, RSPH1*) (3-5). Moreover, little is known about clinical features of patients with different variants in the same PCD gene. Mechanistic explanations for phenotypic differences among patients with the variants in the same gene may lie in modifier genes, microbiome, and environmental factors, among others. Airway inflammation and infection are dominant features of the PCD airway, which may be exacerbated by environmental exposures (e.g., cigarette smoke or allergens) (6). Phenotypic variations are well founded in cystic fibrosis, another genetic cause of chronic airway infection, where disease phenotype can be predicted by the class of mutation in the *CFTR* gene. Nevertheless, individuals with the most common *CFTR* mutation, the homozygous *F508del* variant, have a range of severity of disease, attributed in part to non-*CFTR* modifier genes (7, 8).

Compared to CF, PCD is less common and more genetically heterogenous, which leads to significant challenges in performing genotype-phenotype studies. With few exceptions, individuals with PCD have two pathogenic alleles within the same gene locus as autosomal homozygous recessive (9). Each of the variants may be identical or unique, leading to allele heterogeneity. Predicting the clinical phenotype is particularly difficult in the later condition. Compared to two identical variant alleles, if each allele of the same gene harbors a different mutation, the effect of residual function or total absence of the protein may ultimately dictate cilia function. This allele dosage affect has not been tested in PCD.

To test the contribution of gene dosage to genotype-phenotype relationships in PCD, we investigated mutations in the dynein axonemal assembly factor *DNAAF5* (*HEATR2*), a gene causative of PCD (10). DNAAF5 is a member of the HEAT-repeat-containing family of proteins (*H*untingtin, *e*longation factor 3, PP2*A*, m*T*OR) (11, 12) characterized by ∼30-40 amino acid long modules within alpha helical repeats (13). We previously reported that DNAAF5 belongs to a group of at least 11 cytoplasmic proteins called dynein axonemal assembly factors (DNAAFs) (10, 14, 15). DNAAFs, including DNAAF5, are expressed only in the cytoplasm, are not found in the cilia, and are responsible for the assembly of the components of the ciliary motors within the cytoplasm, prior to transport to the cilia. Pathologic variants of any of the DNAAFs result in absence of the large motor protein complexes within the ciliary axoneme, called outer and inner dynein arms (**Supplemental Figure 1A**), which result in ciliary dysmotility (10, 15-21). Unlike many of the PCD associated proteins, which have variants throughout the cDNA sequence and evidence of deletions/nulls variants, *DNAAF5* is unique as only missense and deletion variants have been identified, and no evidence of homozygous human null variants were reported in ClinVar (22).

We previously described individuals with different *DNAAF5* variants, their spectrum of clinical disease, and their abnormal protein products (14). We leveraged our patients’ clinical features and associated variants in *DNAAF5* to develop mouse models harboring these mutations. Modeling specific human mutations in mice has been a powerful tool for understanding disease pathobiology while controlling for genetic background and environment (23-25). Models of human PCD variants in mice have not been reported. Knockout (KO) of genes causative of PCD in mice have a characteristic phenotype featuring sinus but not lung disease, infertility, and are dominated by early hydrocephalus and death within about two months. To date, hydrocephalus provides the most robust marker of motile cilia dysfunction in mice (26). To investigate *DNAAF5* variants phenotypes, we used genome editing to introduce a PCD patient missense variant in a conserved region of mouse *Dnaaf5*, as well as a frameshift deletion resulting in a null allele. We found that different pathogenic mutations in the same gene result in diverse phenotypes, similar to patterns observed in patients (14). *Dnaaf5* missense variants demonstrated a gene dosage effect with different allele numbers and combinations resulting in a range of disease phenotypes. Motile ciliated cells in some tissues were also differentially sensitive to the same mutations in *Dnaaf5*; sperm flagella being more affected than airway and Fallopian tube cilia. Finally, proteomic, and transcriptional analysis suggested a broader effect of variants than previously appreciated.

## Results

### Spectrum of PCD phenotypes in patients with pathologic variants in DNAAF5

We have identified several variants in *DNAAF5* as causative of PCD in our patient population (**Supplemental Figure 1B, Supplemental Table 1**). Most of the reported missense or deletion homozygous pathogenic variants fall within the highly conserved HEAT-repeat 10 at the C-terminus of the protein (10, 22, 27). There are reported pathogenic variants in other regions of *DNAAF5* and some variants are of unknown significance (**Supplemental Table 1**). Homozygosity for a *DNAAF5* null allele has not been reported and is not present in the patient population cared for in our centers (>15 patients). The Genome Aggregation Database (gnomAD 3.1.2) shows no homozygotes for any predicted loss of function variants in DNAAF5 (22). Also, gnomAD SVs 2.1 shows no structure variants (SVs) affecting the coding gene itself (28). Furthermore, DECIPHER (a genotype and phenotype database) shows no loss of function sequence variants, but lists 93 copy number variants (CNVs) and 6 “other” variants (such as uniparental disomy), all of which are present as heterozygous (29). In addition to a reported case (30), we identified a patient with compound heterozygosity of a null and missense alleles (**Supplemental Table 1**). These genomic data strongly support that *DNAAF5* is an essential human gene, and PCD patients must have some functional, albeit significantly reduced, DNAAF5.

**Table 1.** Effect of *Dnaaf5* variant gene dosage on cilia phenotype.

| Genotype | Alleles | Survival | CBF<br>(airway) | Sperm<br>motility | Hydro-<br>cephalus | Cilia US<br>(airway,<br>ependyma) |  |
| --- | --- | --- | --- | --- | --- | --- | --- |
|  |  |  |  |  |  | ODA | IDA |
| *Dnaaf5*<sup>WT/WT</sup> | 2 WT:<br>WT/WT | NL | NL | NL | None | NL | NL |
| **NULL allele** |  |  |  |  |  |  |  |
| *Dnaaf5*<sup>WT/NULL</sup> | 1 WT:<br>WT/NULL | NL | NL | NL | None | NL | NL |
| *Dnaaf5*<sup>NULL/NULL</sup> | 0 WT:<br>NULL/NULL | Embry-<br>onic<br>lethal | - | - | - | - | - |
| **MIS allele** |  |  |  |  |  |  |  |
| *Dnaaf5*<sup>WT/MIS</sup> | 1 WT + 1 MIS:<br>WT/MIS | 100% at<br>100 d | NL | NL | None | NL | NL |
| *Dnaaf5*<sup>MIS/MIS</sup> | 2 MIS:<br>MIS/MIS | ~50% at<br>100 d | Dec | Absent | Frequent | ↓ | ↓↓ |
| **Compound**<br>**heterozygous** |  |  |  |  |  |  |  |
| *Dnaaf5*<sup>MIS/NULL</sup> | 1 MIS:<br>MIS/NULL | 0% at<br>100 d | Absent | Not<br>done | All | ↓↓ | ↓↓ |

Abbreviations: CBF, cilia beat frequency; Dec, decreased; MIS, Missense variant, MIS; NL, Normal; US, Ultrastructure; WT, Wild type. Arrows: ↓, decreased; ↓↓, very decreased

Indeed, our *DNAAF5* cohort (>15) have a range of clinical features, including typical symptoms of PCD: chronic cough, nasal symptoms, and recurrent otitis media. Organ laterality changes (including *situs inversus* and congenital heart disease) are present in ∼50% of patients, as predicted. When measured, nasal nitric oxide levels are uniformly low, consistent with PCD (31). Transmission electron microscopy (TEM) of *DNAAF5* variants typically show absence or truncation of the outer dynein arms, and absence of the inner dynein arms, thought to be due to a failure of the motor complex to assemble in the cytoplasm, then move to the axoneme microtubules (14, 15, 19). Patients live in diverse environments including urban settings and a cohort in a Mennonite farming community. These exposures could influence phenotypes (32).

Intriguingly, we identified a family with a *DNAAF5* homozygous missense variant (c.14999G), resulting in the substitution of cysteine by phenylalanine at position 500 (C500F) (14). Affected siblings show typical diagnostic features of PCD including *situs* abnormalities, low nasal nitric oxide, and reduced numbers of inner and outer dynein arms determined by TEM. However, compared to other patients with *DNAAF5* variants, these children have less severe disease, and normal cilia beat frequency was retained in about a quarter of the cells, as quantified in cultured nasal epithelium (14). Furthermore, while the numbers of inner and outer dynein arms were reduced, the extent of reduction was less pronounced than in patients with other *DNAAF5* variants (**Figure 1A and 1B**). Given the lack of reports of individuals with biallelic *DNAAF5* null alleles, as well as the existence of patients with clinically milder PCD features, we asked whether *DNAAF5* variants represented an allelic series, or whether the clinical variation could be attributed to environment or the presence of modifier genes.

**Figure 1.**
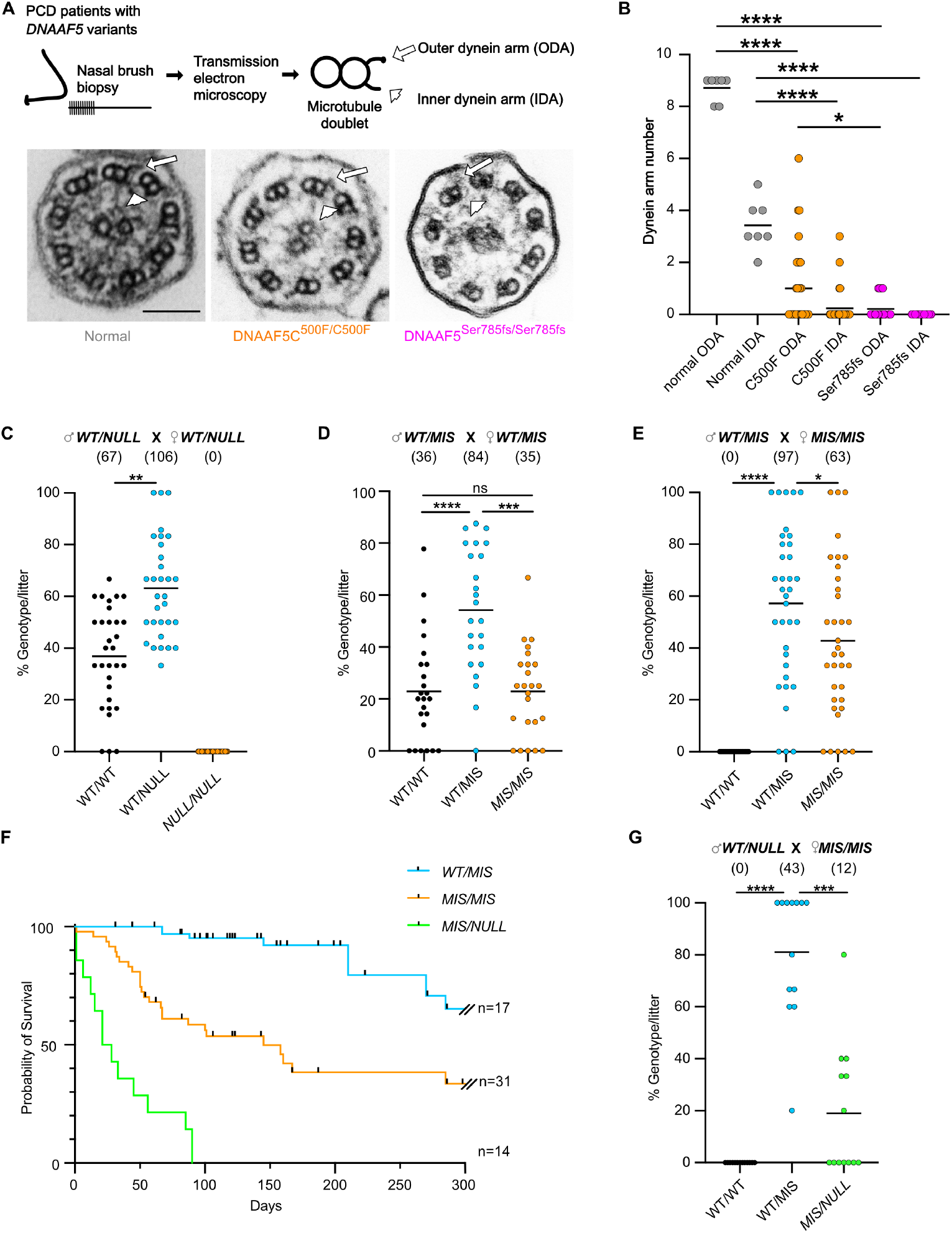
The effect of NULL and MIS allele dose on survival. **(A**) Nasal epithelial cells were obtained from patients with *DNAAF5* variants for transmission electron microscopy (TEM) to assess DNAAF5 function by the ultrastructure of the ciliary axoneme. Cross section of cilia show normal, truncated, or absent outer (ODA) and inner (IDA) dynein arms (cilia motor complexes) on microtubule doublets. Arrow and arrowhead indicate the ODA and IDA, respectively. Bar=100 nm. (**B**) Quantification of ODA and IDA identified in TEM cross sections of cilia from individuals with indicated variants in *DNAAF5* (n=1-2 individuals per genotype). (**C**) Intercross of *WT/NULL* mice produce no *NULL/NULL* offspring (n=31 litters, 173 mice were analyzed). Mean±S.E. percent of genotype/litter: 36.9±19.3%, 63.2%±19.3%, 0% in *WT/WT, WT/NULL, and NULL/NULL, respectively*. (**D**) Intercross of male *WT/MIS* and female *WT/MIS* mice (n=24 litters). Mean±S.E.percent of genotype/litter: 22.9±4.2, 54.2±5.0, and 22.9±3.5 in *WT/WT, WT/MIS, and MIS/MIS, respectively* (**E**) Intercross of *WT/MIS* males with *MIS/MIS* females (n=35 litters, mean=57.2%±5.0 *WT/MIS*, 42.8 ± 5.0, *MIS/MI*). Fewer *MIS/MIS* offspring were produced than predicted by Mendelian genetics (Chi square, p=0.04). (**F**) Kaplan survival curve for mice with indicated genotypes. (**G**) Breeding *WT/NULL* males with *MIS/MIS* females produce fewer than predicted MIS/NULL offspring (n=13 litters, mean=81.0±7.0%, *WT/MIS*, 19.0%± 7.0, *MIS/NULL*; Chi square p=0.0009). Means are shown in B-E and G; *P<0.05, **P<0.01, ***P<0.001, ****P<0.0001 determined using Kruskal-Wallis test with Dunn’s multiple comparisons. Number of animals in each group is shown in parenthesis in C,D,E,G.

### Different mutations in Dnaaf5 result in distinct phenotypes

To investigate the phenotype-genotype relationships in an identical genetic background and environment, we used a CRISPR-Cas9 system to introduce the human c.14999G in the conserved *Dnaaf5* mouse genome at position c.1493, resulting in a G>T and the equivalent p.C498F missense mutation (**Supplemental Figure 2A**). A synonymous mutation (c.1473;C>T) was also introduced upstream to the missense mutation to prevent genomic re-cutting by Cas9 and improve the probability of homology-directed repair (**Supplemental Figure 2A**) (33).

Surrogate mothers implanted with edited blastocysts, all in the C57BL/6 strain, provided chimeric pups. Fifty-two founder pups were born, and genotype determined. Fifteen were wild type (WT) with no disease phenotype, one of the pups showed a heterozygous missense mutation without a difference from the WT phenotype at birth (c.1493, G>T), and 26 had indels resulting in multiple frameshift mutations. Mice with indels had features typical of PCD gene knockout (KO) mice, especially hydrocephalus and runted growth (**Supplemental Figure 2B and 2C**). Surviving mice with indels carried one of three different frameshifts (FS), each in *trans* with a WT allele (**Supplemental Figure 2D**). Germline transmission of the FS and missense C498F *Dnaaf5* alleles were established after breeding with wild-type mice. The heterozygous F1 pups (*Dnaaf5*^*WT/FS*^ and *Dnaaf5*^*WT*/C498F^) had normal post-natal growth and behavior. Germline transmission was confirmed in all three FS lines by breeding. We used the line with a c.1476delGGAGCAT FS* (p.Asp493fs*26) mutation for subsequent phenotypic analyses of the FS mutation. The FS mutation is predicted to result in RNA decay and a null allele. The two alleles, FS and NULL, provided an opportunity to study phenotypes of each allele and the gene dosage effects in recessive genetic models (**Table 1**).

An initial intercross of heterozygous *Dnaaf5*^WT/FS^ mice (denoted *WT/NULL*), yielded 31 litters and 173 animals (**Figure 1C**). However, no animals homozygous for the FS mutation (*Dnaaf5*^FS/FS^; denoted *NULL/NULL*) were present at weaning. Lack of homozygous FS offspring was further confirmed by interbreeding the other two frameshift mutation lines. These observations suggest that loss of *Dnaaf5* results in embryonic lethality. To determine when *Dnaaf5* mutants die, independent editing of the critical mouse exon to generate a null allele was performed (**Supplemental Figure 3A**). In this line, Mendelian ratios at blastocyst stage were normal (**Supplemental Figure 3B**). However, by E6.5 (E, embryonic day) homozygous mutant embryos had growth arrest and reduced numbers, and by E8.5 no surviving mutants were observed (**Supplemental Figure 3B and 3C)**. This early embryonic lethality, at a window before cilia function is required, suggests a non-ciliary requirement for DNAAF5. The findings align with observations in *Drosophila* of adult lethality of *CG31320/DNAAF5* mutant flies, where motile cilia are restricted to the sperm and chordotonal neuron (CH) in the fly, and motile cilia defects are not lethal (27, 34). The exact role of the DNAAF5 protein in non-ciliary functions is not clear but is supported by the broad DNAAF5 expression beyond motile ciliated tissues (35). Consistent with data in gnomAD and ClinVar that no *DNAAF5* homozygous null variants are reported, our observed early embryonic lethality in multiple null alleles suggest that *DNAAF5* is an essential Metazoan gene.

Next, to assess the genetic fitness of the other *Dnaaf5* mutations, we interbred *Dnaaf5*^*WT*/C498F^ (*WT/MIS*) mice; the live offspring included all predicted genotypes (**Figure 1D**). The genotype distribution was near Mendelian: 23:54:23%, for wild-type, heterozygotes, and homozygotes, respectively. Breeding male *WT/MIS* with female *MIS/MIS* resulted in live *MIS/MIS* offspring, but less than predicted, which may be related to *in utero* loss of *MIS/MIS* animals or subfertility of WT/MIS females (**Figure 1E**).

*MIS/MIS* animals showed gross phenotypes associated with motile cilia deficiency phenotypes commonly observed in mice, including development of hydrocephalus and *situs* abnormalities (**Supplemental Figure 3D and 3E**). Most animals with two *MIS* alleles (*MIS/MIS*) survived beyond 150 days of life (a subset of animals was followed for up to 1 year) (**Figure 1F**). These findings suggest that homozygous missense mutants retain sufficient function to survive, in contrast to the early embryonic lethality of null alleles.

To study how dosage of a single *MIS* allele affects survival, we bred *MIS/MIS* female animals to *WT/NULL* male animals. Breeding resulted in viable litters (**Figure 1G**). However, only 22% of offspring were *MIS/NULL* (predicted 50%), suggesting *in utero* mortality of most *MIS/NULL* offspring. Compound mutant *MIS/NULL* mice that were born alive, were runted and displayed early onset gross macrocephaly (**Supplemental Figure 3F and 3G**). Early lethality was observed with 50% of these compound mutant mice dying by 30 days and none surviving beyond 90 days (**Figure 1F**).

Taken together, our results indicate that *Dnaaf5* is essential for development and sensitive to gene dosage. While complete loss of function of *Dnaaf5* (*NULL/NULL*) is embryonic lethal and homozygous missense C498F (*MIS/MIS*) survive postnatally, compound mutants (*MIS/NULL*) exhibit increased perinatal lethality, suggesting we have generated mice representing an allelic series of *Dnaaf5* function, *in vivo*.

### Combinations of Dnaaf5 alleles determine cilia motility and transport

Both our patient phenotype and our *MIS* allele analysis suggest that missense mutant protein may be partially functional, with two *MIS* alleles providing an additive effect. To test the effect of allelic dosage on cilia phenotype, we evaluated the cilia function by measuring cilia beat frequency (CBF) and cilia transport of beads in the trachea of mice *ex vivo* (**Figure 2A**). In parallel, we also cultured primary tracheal epithelial cells at air-liquid interface (ALI) to exclude environmental factors (e.g., infection) (36).

**Figure 2.**
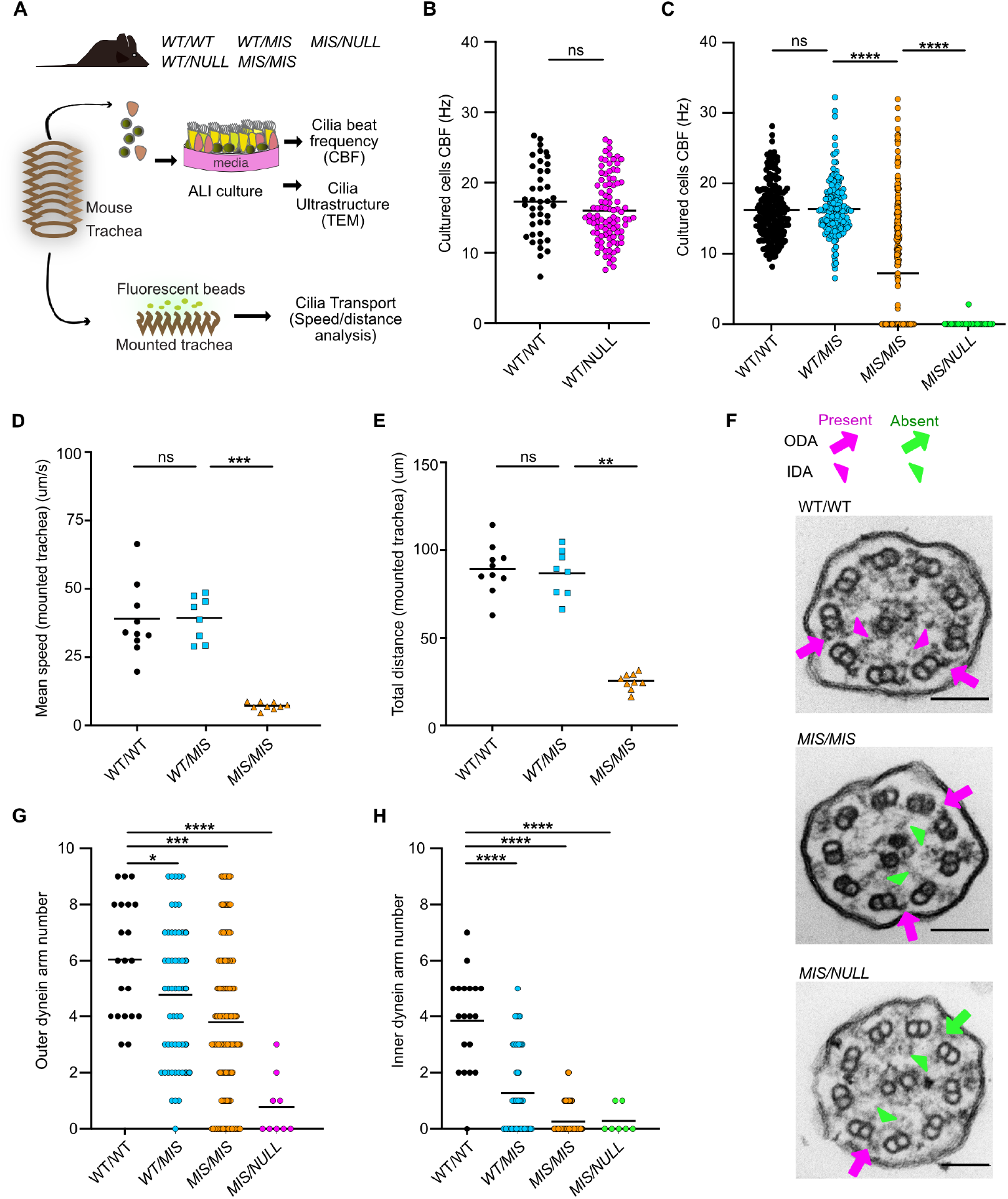
The effect of *NULL* and *MIS* allele dose on airway cilia function and ultrastructure. (**A**) Schematic of primary culture airway cells and ex vivo isolated trachea mounts used for analysis of cilia function by cilia beat frequency (CBF), transport and ultrastructure by transmission electron microscopy (TEM). (**B**) CBF of cultured cells from *WT/WT* and *WT/NULL* littermates. Mean CBF 17.47±0.80 and 16.0±0.47 Hz, respectively. (**C**) CBF of cultured cells from *WT/WT, WT/MIS, MIS/MIS*, and *MIS/NULL* littermates. Mean frequency 16.22±0.29, 16.39±0.33, 7.25±0.53, and 0.06±0.06 Hz, respectively. CBF was assessed in 5 random areas per Transwell; 5 Transwell cultures were performed for each genotype per experiment, n=2-4 independent experiments comprising unique mice. (**D**,**E**) Mucociliary transport speed and bead displacement on the surface of isolated trachea from *WT/WT, WT/MIS*, and *MIS/MIS* mice. Mean speeds 39.04±5.02, 39.29± 2.86, and 7.16±0.45 m/sec, respectively. Mean bead displacement over 5 seconds, 89.13±4.42 µm, 86.82±4.80 µm, 26.00±1.36 µm, respectively (n=8-10 animals per genotype). (**F**) Representative TEM images of airway cilia from *WT/WT, MIS/MIS*, and *MIS/NULL* mice. Arrows indicate presence (magenta) or absence (green) of outer and inner dynein arms (ODA, IDA). Bar=100 nm. (**G**) Quantitation of outer dynein motor protein complexes detected by TEM of *WT/WT, MIS/MIS*, and *MIS/NULL* cilia cross section. Mean ODA 6.0±0.4, 3.8±0.2, and 0.8±0.4, respectively. (**H**) Quantification of inner dynein arms in ciliary axonemes from cultured tracheal epithelial cells from *WT/WT, MIS/MIS, and MIS/NULL*. Mean IDA 3.8±04, 0.3±0.0, and 0.3±0.2, respectively. (n=4 animals per genotype, n=5-10 cross sections per genotype). *P<0.05, **P<0.01, ***P<0.001 determined using Kruskal-Wallis test with Dunn’s multiple comparisons.

We first compared the CBF of *WT/WT* to *WT/NULL* littermates, finding no difference in the CBF between these genotypes (**Figure 2B**). Next, we compared the effect of *MIS* dosage on CBF. The effect of a single *MIS* allele when in *trans* with a *WT* allele (*WT/MIS*) was not statistically different from *WT/WT* (**Figure 2C**). Similarly, bead transport across the surface of the trachea was not different between mice of these genotypes (**Figure 2D and 2E**). In contrast, two *MIS* alleles (*MIS/MIS*) resulted in variable but distinctly lower mean CBF and decreased tracheal bead transport when compared to *WT/MIS* and *WT/WT* control littermates (**Figure 2C-2E**). The heterogeneity of cilia motility among cells in the *MIS/MIS* samples was similar to that observed in our patients with the *DNAAF5* c.14999G (*C500F/C550F*) missense variant (14); most cilia from *MIS/MIS* animals were completely immotile, while others retained markedly reduced motility (Supplemental **Video 1-3**).

Finally, we tested a single copy of the *MIS* mutation in *trans* with the *NULL* allele (*MIS/NULL*). *MIS/NULL* cultured cells showed a more pronounced cilia motility defect compared to *MIS/MIS*, as all cilia were immotile (**Figure 2C**). Together, these results suggest that a single copy of C498F results in a Dnaaf5 protein with either reduced levels or functions that are required to adequately equip motile cilia, likely due to inadequate dynein motor complex assembly.

### Dnaaf5 mutations determine the degree of cilia ultrastructure defects

Dnaaf5 contributes to the cytoplasmic assembly of the axonemal inner and outer dynein arms motor complexes (IDA, ODA). The absence of ODA and IDA in the cilia indicates insufficient production of the motor complexes. To investigate the relationship between the CBF phenotype and genotypes, we examined the cilia ultrastructure using transmission electron microscopy.

Cilia were scored for normal or defective (absent, truncated) IDA and ODA, compared to *WT/WT* mice. We did not appreciate structural defects in cross sections of *WT/NULL* cilia compared to *WT/WT*. Consistent with our ciliary beat frequency results, normal axonemal ultrastructure suggests that defects in cilia motility do not arise from haploinsufficiency, i.e., one *WT* allele is sufficient for normal dynein motor assembly. Next, we evaluated the effects of levels of the *MIS* mutation on airway cilia ultrastructure. We observed a dose-dependent reduction in the mean number of ODAs and IDAs with successive loss of the *WT* allele (*WT/WT* to *WT/MIS* to *MIS/MIS*) (**Figure 2F-2H**). IDA were significantly reduced in cross sections of cilia from *MIS/MIS*. To assess *MIS* dosage, the ultrastructure of cilia from mice with a single *MIS* allele was assessed in isolation (*MIS/NULL*). Consistent with the reduced CBF, ODAs and IDAs were also mostly absent in the setting of a single mutant allele (**Figure 2G and 2F**).

As assessed by TEM, there was striking variability in the presence and morphology of ODA and IDA, among different doublet microtubules in the same cilia cross-section of mice with a mutant allele (*MIS/MIS* or *MIS/NULL*). The ODA appeared normal on some microtubules while absent or truncated on other microtubules within the same cross section (**Figure 2F**). This variability was not unique to a specific microtubule of the nine different A tubules (diagram, **Supplemental Figure 1A**) that dock the dynein motor complexes. Variable ODA morphology was similarly present in patients with the *C500F* mutation (**Figure 1A**). This observation may suggest that delivery of the dynein complex is not uniform along the length of the ciliary axonemal and that partially formed complexes may be transported into the same ciliary axoneme alongside fully assembled dynein complexes.

### Tissue specificity of genotype-phenotype relationships: airway

Having observed differences in cilia ultrastructure in the airway, we examined the tissue specificity of phenotypes in mice with different *Dnaaf5* genotypes. Motile cilia dysfunction is associated with upper airway sinus disease in patients and mice with knockout of PCD gene (37-41). Unlike humans, mice deficient in PCD genes are not reported to develop lung disease. We did not observe lung inflammation in our mice of any genotype. We observed sinus inflammation in the *MIS/MIS* and *MIS/NULL* mice, compared to *WT/MIS* mice and WT/WT type-littermates (**Supplemental Figure 4A**). We took advantage of the long survival of *MIS/MIS* mice to test lung clearance using intratracheal *Pseudomonas aeruginosa*. There was a trend toward decreased clearance in the *MIS/MIS* mice compared to *WT/MIS and WT/WT* animals (**Supplemental Figure 4B and 4C**). We were unable to perform these experiments in *MIS/NULL* animals due to limited survival beyond weaning age.

### Tissue specificity of genotype-phenotype relationships: ependyma

Motile cilia are considered essential for creation of CSF flow networks (42) and defects in cilia-related genes are associated with enlarged brain ventricles, i.e., hydrocephalus. Mice deficient in genes coding for dynein assembly factors or dynein arm complex proteins (e.g., *Dnah5*), develop severe postnatal macrocephaly due to hydrocephalus, which is fatal within approximately a month (17, 41, 43, 44). We did not observe gross hydrocephalus in *WT/WT, WT/MIS*, or *WT/NULL* mice. In contrast, about half of *MIS/MIS* and all *MIS/NULL* animals developed at least mild hydrocephalus by 12 weeks of age, ultimately leading to death. However, in animals surviving longer (1 to 10 months), histologic and brain magnetic resonance imaging (MRI) showed varying degrees of ventriculomegaly in all *MIS/MIS* animals, independent of the presence of gross macrocephaly or sex (**Figure 3A and 3B)**. By magnetic resonance imaging (MRI), the lateral and third ventricle in *MIS/MIS* animals were significantly larger than in *WT/MIS* and *WT/WT* animals (**Figure 3C and Supplemental Figure 5A**), while there was no difference in fourth ventricle volumes (**Supplemental Figure 5B and 5C**). Histologic evaluation did not show obstruction within the brain ventricle system or aqueduct, suggesting that *MIS/MIS* and *MIS/NULL* animals develop non-obstructive, communicating hydrocephalus due to insufficient DNAAF5 function leading to insufficient motor complexes in the cilia.

**Figure 3.**
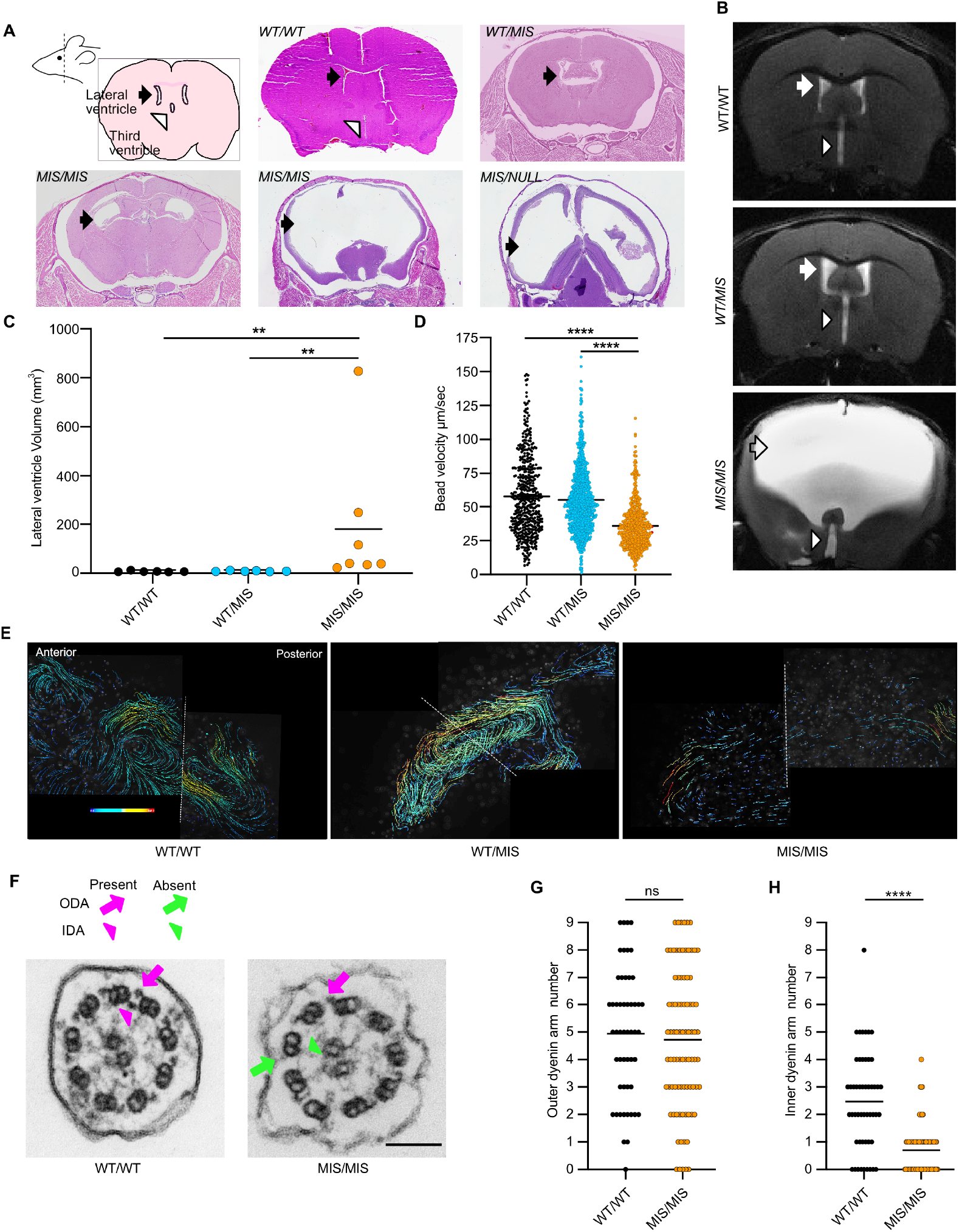
The effect of *MIS* allele dose on hydrocephalus and ependymal cilia ultrastructure. (**A**) Coronal brain cross sections show hydrocephalus in *MIS/MIS* and *MIS/NULL* animals compared to other genotypes. Note variable degrees of hydrocephalus in *MIS/MIS* mice. Arrow indicates lateral ventricles. (**B**) MRI images showing ventricular enlargement in *MIS/MIS* compared to *WT/MIS* and *WT/WT* animals. Arrow indicates lateral ventricles; arrowhead indicates third ventricle. (**C**) Quantification of lateral ventricle volumes from MRI images in C showing significant ventricular enlargement of *MIS/MIS* compared to *WT/MIS* and *WT/WT* mice (P<0.01, n=6 animals per genotype). (**D**) Bead velocity on isolated ventricles from *WT/WT, WT/MIS*, and *MIS/MIS* mice. Mean speeds were 57.57±1.2, 55.02± 0.78, and 35.93± 0.70 µm/sec, respectively, and were significantly different between *WT/WT* and *MIS/MIS* animals (P<0.0001, n=2 animals per genotype, measured on 2 lateral ventricles per animal). (**E**) Representative composite images of fluorescent beads velocity vectors showing transport on ex vivo lateral ventricles. (**F**) Representative TEM images of cross sections from ependymal cilia of *MIS/MIS* and *WT/WT* littermates showing reduced dynein motor protein complexes in ependyma ciliary axonemes of *MIS/MIS* mice. Arrows and arrowhead indicate the presence (magenta) and absence (green) of ODA and IDA, respectively. Bar=100 nm. (**G-H**) Quantification of ODA and IDA in ependymal cilia cross sections from different genotypes (n=2 animals per group). *P<0.05, **P<0.01, ***P<0.001 determined using Kruskal-Wallis test with Dunn’s multiple comparisons.

In addition to brain abnormalities, *MIS/MIS* mice with hydrocephalus had cervical spine lordosis with thoraco-lumbar kyphosis, detected by MRI (**Supplemental Figure 5D**). Even *MIS/MIS* mice with mild ventricular enlargement had the same spinal abnormality observed in mice with severe ventricle enlargement. No skeletal deformity was noted, suggesting that the spine deformity may be secondary to neuromuscular changes. This spine phenomenon was observed in other mice that are null for motile cilia genes, including *Foxj1*^-/-^ animals (43, 45). In zebrafish, spinal curvature is often observed in motile ciliopathy and is linked to dysfunction of the subcommissural organ and Reissner’s fiber within the ventricular system (46, 47).

To examine the effect of *Dnaaf5* mutant alleles in the cilia of the ependyma compared to the airway, we performed transport assays on the surface of extracted lateral ventricles of *WT/WT, WT/MIS*, and *MIS/MIS* animals (**Figure 3D**). *MIS/NULL* were not evaluated as severe hydrocephalus limited extraction of the brain with intact ventricles. Cerebral spinal flow is directed by complex patterns of cilia-mediated transport (42). The rate and distance of movement of beads on the surface of ependymal cilia on the lateral ventricles of *WT/WT* and *WT/MIS* mice were similar, in contrast *MIS/MIS* animals had comparatively slower bead velocity and shorter flow distance (**Figure 3D and 3E;** Supplemental **Video 4 and 5**). The loss of ependyma ciliary axoneme IDA and ODA in *MIS/MIS* compared *WT/WT* were similar to those observed in airway cilia (**Figure 3G and 3H**).

Taken together, these observations suggest a dose-dependent effect of the *MIS* allele on development of hydrocephalus. In the presence of a single *C498F* allele (*MIS/NULL*), hydrocephalus occurred earlier and was more severe than in *MIS/MIS* animals.

### Tissue specificity of genotype-phenotype relationships: fertility

Fertility is related to motile cilia in the Fallopian tube and the function of the sperm flagella. We first tested the effect of gene dosage of the *MIS* allele in female mice by breeding *Dnaaf5 WT/MIS* or *MIS/MIS* females with *WT/WT* males, which resulted in live offspring indicate that *MIS/MIS* female mice are fertile. To assess sperm function, *WT/MIS* or *MIS/MIS* males were bred with *WT/WT* females. Only *WT/MIS* males produced live offspring, indicating that two missense alleles in *Dnaaf5* cause male infertility. Limited survival prevented testing the fertility of *MIS/NULL* mice of either sex. These findings suggest that the sperm are more sensitive to mutations in *Dnaaf5* than the Fallopian tube cilia.

To assess the effects of these mutations on sperm and Fallopian tube cilia function, we evaluated the cilia motility in the relevant tissues. Unlike in the airway and ependyma cilia, where decreased motility was observed in *MIS/MIS* cilia, sperm from *MIS/MIS* animals were all immotile and could not be recovered by capacitation or the addition of 8-Bromo-cAMP, despite similar sperm viability between genotypes (**Figure 4A and 4B, Supplemental Figure 6A and Supplemental Videos 6 - 10)**. There were no significant differences in sperm motility between *WT/WT* and *WT/MIS* mice.

**Figure 4.**
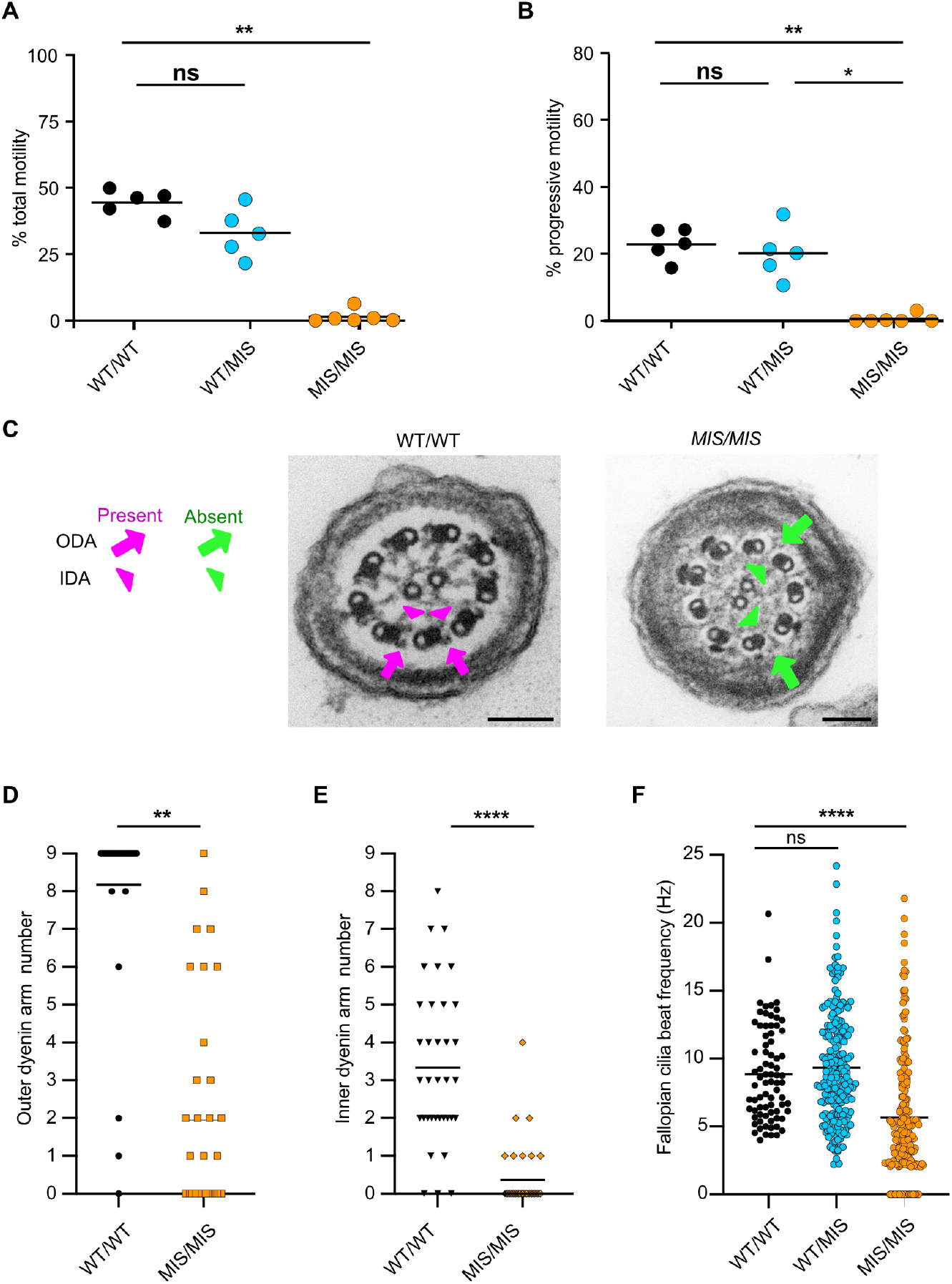
The effect of *MIS* allele dose on sperm and Fallopian tube function and structure. (**A**) Sperm with motility was low in *MIS/MIS* (1.35±1.0%) compared to *WT/WT* (44.46±2.17%) and *WT/MIS* (33.04±4.10%) littermates (n=5-6 mice/genotype, P<0.01). (**B**) Sperm showing forward progressive motility was significantly lower in *MIS/MIS* sperm (0.56±0.5%) compared to *WT/WT* (22.90±2.11%) and *WT/MIS* (20.12±3.27%) littermates; 0.56±0.5%, 22.90±2.11%, and 20.12±3.27%, respectively (n=5-6 mice per genotype. P<0.01). (**C**) Reduced dynein arms in *MIS/MIS* sperm compared to *WT/WT* littermates. Representative TEM images of cross sections from sperm flagella. Arrows and arrowhead indicate the presence (magenta) and absence (green) of ODA and IDA, respectively. Bar=100 nm. (**D-E**) Reduction of motor complex ODA and IDA in sperm flagella in *MIS/MIS* compared to *WT/WT* littermates (n=2 mice per genotype). (**F**) Reduce CBF of cilia from ex vivo, isolated Fallopian tube from *WT/WT, WT/MIS, and MIS/MIS* littermates, 5.65±0.31, 8.85±0.39, and 9.34±0.28 Hz, respectively (n=2-3 mice per genotype). *P<0.05, **P<0.01, ****P<0.0001 determined using Kruskal-Wallis test with Dunn’s multiple comparisons.

Consistent with loss of function, ultrastructural evaluation of flagella of *MIS/MIS* sperm showed complete absence of outer and inner dynein arms (**Figure 4C-4E**). Interestingly, approximately half of the sperm cross-sections from *MIS/MIS* animals also showed abnormal numbers of doublet microtubules and absence of the central apparatus (**Supplemental Figure 6B and 6C**). This ultrastructural abnormality was not observed in airway cilia from either mice or patients with mutations in *Dnaaf5* and may be specific to sperm.

We also assessed Fallopian tube cilia function. Unlike male *Dnaaf5 MIS/MIS* animals, female animals with *MIS/MIS* mutations were fertile. Indeed, analysis of cilia beat frequency of dissected Fallopian tubes showed motile cilia, with a frequency range similar to airway cilia (**Figure 4F**).

In summary, differences in cilia function and accompanying ultrastructural changes in sperm compared to airway suggest differences in the tolerance of mutations in dynein motor assembly proteins in different tissues.

### Ciliary axoneme proteomics of Dnaaf5 mutant mice

How *Dnaaf5* variants cause different phenotypes could be related to the quantities or types of motor proteins missing in the axoneme of mutants. Cilia from patients with PCD variants in genes coding for DNAAFs have absent ODA and IDA ciliary dynein arm proteins, including DNAH5, DNAI1, DNALI1, and DNAH7, in ciliary axonemes, however an unbiased analysis of proteins in the variant ciliary axoneme has not been performed (10, 27). We isolated cilia from cultured tracheal epithelial cells from mice with the *MIS/NULL* genotype and compared proteomics to wild-type littermates by tandem mass tag (TMT) mass spectrometry (**Figure 5B and Supplemental Table 2**).

**Figure 5.**
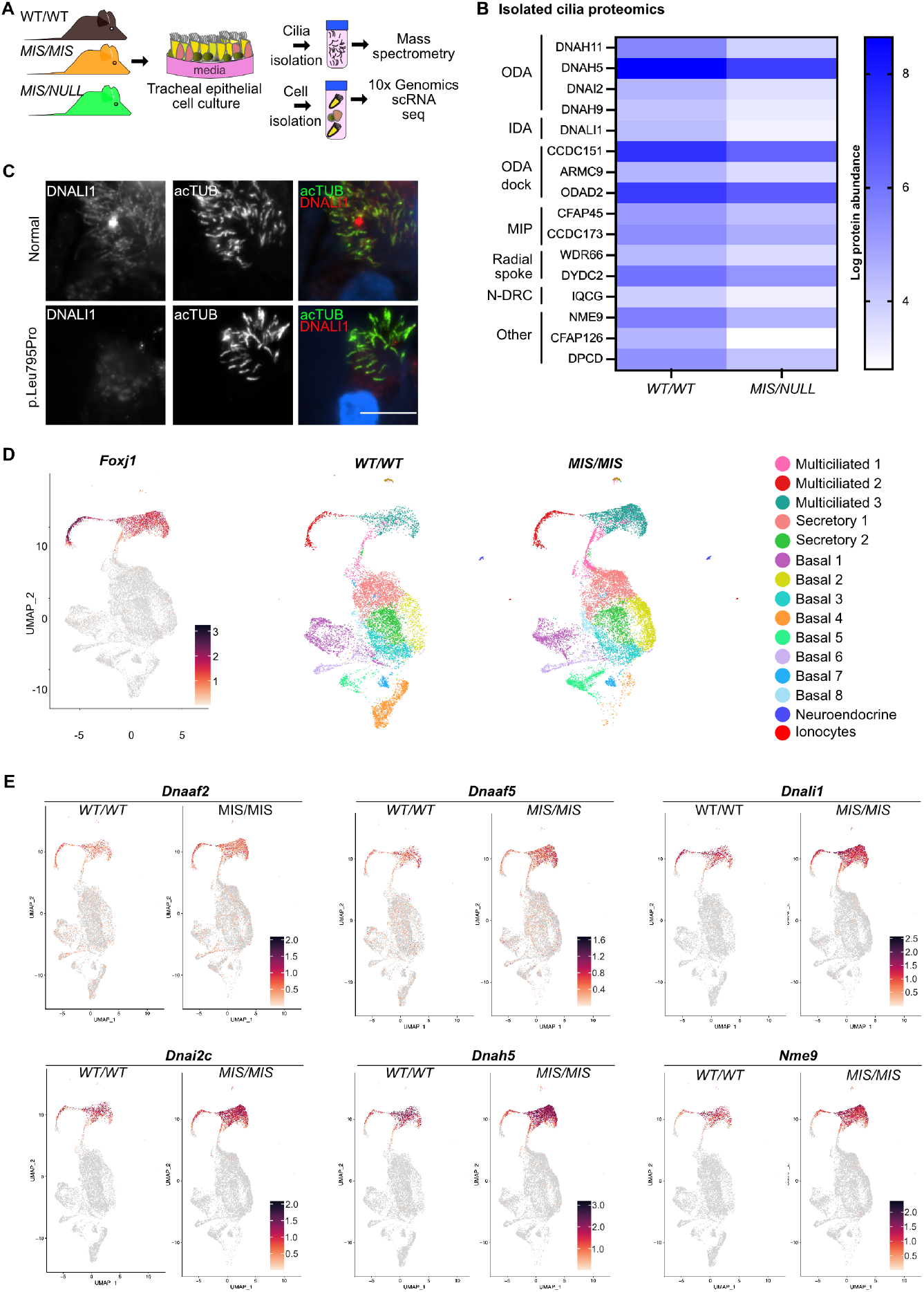
Molecular phenotype of *Dnaaf5* mutant multiciliated cells. (**A**) Schematic of strategy used for analysis of cultured airway cells from wild type and mutant mice. Isolated motile cilia were analyzed by mass spectrometry and total cells by single cell RNA sequencing. (**B**) Proteomics of ciliary axonemes isolated from primary culture tracheal epithelial cells from *WT/WT* and *MIS/NULL* littermates. Heat-map comparing the expression level of cilia-associated proteins (n=4 animals per genotype). (**C**) Reduction in DNALI1 in cilia of primary human airway cells from a normal individual and a patient with PCD due to *DNAAF5* variant p.Leu795Pro by immunofluorescent analysis. Representative images shown, bar=100 µm. (**D**) UMAP comparing airway epithelial cell clusters of *WT/WT* and *MIS/MIS* littermates. Foxj1 expression is shown for reference (n=3 animals per genotype). (**E**) Feature-plot showing differential expression of selected genes in *WT/WT* and *MIS/MIS* littermates identified using scRNAseq analysis.

Compared to *WT/WT, MISS/NULL* axonemal cilia showed significant reduction in ODA proteins, DNAH11, DNAH5, DNAI2, and IDA protein, DNALI1 (**Figure 5B**). Interestingly, we also observed reduction in proteins associated with the ODA docking complex, including CCCDC151 and ARMC4, suggesting that the docking proteins are interdependent on motor protein assembly, transport or retention in the axoneme. Proteins in the cilia with less known function were also identified as reduced, include CFAP126 (Flattop), a microtubule inner protein, and NME9, a thioredoxin domain-containing protein (**Figure 5B and Supplemental Table 2**). Analysis of primary airway cells from our patients with a pathogenic variant in *DNAAF5* showed decreased expression of some of these candidate proteins in the ciliary axonemes (**Figure 5B**, and **Supplemental Figure 7A**). These missing proteins were not previously associated with *Dnaaf5* mutations, which may suggest a wider disruption of the integrity of dynein arm complexes, or other roles for DNAAF5 (and other DNAAF proteins) in ciliary motor assembly and cilia assembly.

### Dnaaf5 mutation are associated with increased cilia-related transcript abundance

The effect of mutations in cilia assembly genes on the airway multiciliated cell transcriptome is not defined. We hypothesized that mutations in *Dnaaf5* result in a compensatory response to decreased motor proteins in the ciliary axoneme. We used single cell RNA sequencing to transcriptionally profile cultured tracheal epithelial cells from *Dnaaf5 MIS/MIS* and wild-type littermates (n=3 unique mice for each genotype) (**Figure 5C**). We identified all known major cell epithelial types, including basal progenitor cells, secretory cells, multiciliated cells, ionocytes and neuroendocrine cells (**Supplemental Figure 7B and 7C**). *MIS/MIS* mutations did not affect the distribution of cell types or differentiation of culture primary cells compared to *WT/WT*, though we did observe a shift in the abundance of subclusters of basal cells between groups (**Figure 5D)**. Ontology analysis showed increased expression of cilia assembly as a group (GO: 0060271), with *MIS/MIS* cells showing increased differential expression of motile cilia related genes (e.g., transcripts of outer dynein arm genes DNAH5, radial spoke gene RSPH9, and organization genes CCDC39 **(Figure 5E, Supplemental Figure 7D and Supplemental Table 3)**. Downregulated transcripts in motile ciliated cells were related to glutathione metabolism and prostaglandin metabolism (**Supplemental Table 4**). To determine if the same molecular phenotype was present in patients with *DNAAF5* variants, we examined bulk RNA from primary culture nasal epithelial cells obtained from a patient with a compound variant (*DNAAF5*, c.2353-2354del,p.S785 fs). The upregulated genes were similar to those observed in *Dnaaf5* mutant mice cells (**Supplemental Figure 7E**), suggesting a conserved feedback response in human disease, broadly affecting ciliogenesis.

## Discussion

Patients with PCD have pathologic allele variants leading to clinical features affecting specific organ systems: the respiratory tract, reproduction, cerebral spinal fluid circulation and development of organ laterality. Yet, the disease of PCD is not invariable, instead patients have a constellation of signs and symptoms that differ among individuals, across ages, and we propose, by gene variant and mutation type. We also recognize that environmental influences, infection type, and variant suppressor gene may influence phenotypes. To address these issues, we investigated PCD genotype-phenotype relationships in a fixed genetic background and environment by modeling human PCD variants of *Dnaaf5* in mice. We drew upon our observations of patients with *DNAAF5* variants, experience with DNAAF5 function in ciliogenesis, and familiarity with the unique features of motile cilia dysfunction in mice (10, 14, 45). We were particularly interested in a human *DNAAF5* variant associated with mild disease that we previously showed produced detectable protein, determined by immunoblot (14). Generating a clinically conserved allele to model a human missense variant and a null mutation in mice by CRISPR-Cas9 genome editing allowed formal testing of allele fitness and the impact of each allele alone and in combination with a wild type allele. Using this approach, we identified a distinct gene dosage effect of *MIS* and *NULL* alleles (**Table 1**). In mice, gene dosage was associated with clinical phenotypes of motile cilia-dependent organs and quantifiable differences in cilia motor function and cilia ultrastructure, emphasizing the importance of knowledge of the specific mutation, the function of the gene and, if relevant, the allele combinations.

Survival served as a quantifiable measure of allelic fitness. Mice with PCD gene knockout uniformly develop early hydrocephalus and death by two months of age (or earlier) (39-41). While rare in humans with PCD, development of hydrocephalus is a well-established phenotype in mice, which we confirm is consistent with defective function and ultrastructure of cilia on the ventricular ependyma (48). The development of ventriculomegaly followed by an early death is similarly observed in knockout of most motile cilia genes in the hydrocephalus prone C57BL/6 strain, limiting the ability to identify differences in phenotypes (26, 49). In contrast to the knockout approach, we opted to introduce a patient specific mutation (*DNAAF5*^*C500F/C500F*^, *MIS/MIS)* and a null mutation to study the interaction of different variants (14). We first assessed allelic fitness of the *NULL* and *MIS* alleles by their survival. We demonstrate that in the heterozygous condition, in a wild-type background, neither allele (*WT/MIS* or *WT/NULL*) significantly impacted the survival phenotype. The null allele (*NULL/NULL*) in the homozygous condition however resulted in early embryonic lethality, which, to our knowledge, has not been previously noted in PCD gene knock out models. The homozygous missense mutation (*MIS/MIS*) resulted in decreased survival, while in *trans* with a single null allele. the *MIS/NULL* genotype allowed live birth, but death within one month. Thus, when not paired with the *WT* allele, each mutant allele alone was lethal, but to a different degree. These results provide evidence that phenotypes should be interpreted relative to the specific variant, which may impact our analysis of genotype-phenotype relationships in human disease.

The cause of embryonic lethality observed in animals with *NULL/NULL* mutations in *Dnaaf5* is unclear, but given it arises by E6.5, in a window prior to cilia function, earlier than motile cilia function in nodal cilia at E7.5 (50). The results point to a cilia-independent chaperoning function for DNAAF5 during embryogenesis. As noted, the orthologue of *Dnaaf5* is also lethal in early *Drosophlia* development (27). The early intrauterine death of mice suggests that *DNAAF5* variants in humans have some partial function, at least pertaining to the survival phenotype. Indeed, we were able to detect reduced levels of DNAAF5 by immunoblot in cultured nasal epithelial cells from patients with DNAAF5 variants (14). Second, the majority of known *DNAAF5* human deletion variants are within the 3’-terminus (**Supplemental Figure 1 and Supplemental Table 1**), likely retaining a sufficient, partially functioning protein. Third, homozygous null mutations of *DNAAF5* are not found in ClinVar or gnomAD, indicating that pressures of allelic fitness are at work. Additional support for survival of variants with partial function comes from *Chlamydomonas*, where truncated abnormal appearing dynein complex are partially functional, despite the loss of the outer dynein arm motor protein heavy chain beta (oda4) (51).

Fertility was used as a second determinant of allelic fitness. Most males with PCD are infertile, though subfertility occurs and is most often the case in females (52). As expected, male mice with a single copy of the wild-type allele were fertile, consistent with the carrier state in autosomal recessive genetics. Notable was the loss of fertility in *MIS/MIS* male mice, and lack of sperm flagella motility, despite having less frequent hydrocephalus and longer survival than the *MIS/NULL* mice. Observing completely immotile *MIS/MIS* sperm contrasted with decreased retained motility of some cilia in the airway, ependyma and Fallopian tube, likely due to a different motor protein composition of the motor complex in different tissues. The components of the outer dynein motor complex within the axoneme of the airway, ependyma, and Fallopian tube are similar including dynein heavy chains DNAH5, DNAH11, and DNAH9. Sperm dynein heavy chain however differ, using DNAH8 and DNAH17 instead (53). Different cilia gene requirements for flagellar assembly than airway assembly may account for infertility without PCD (54). TEM examination of sperm tail ultrastructure of *MIS/MIS* mice showed significantly diminished outer dynein arms compared to the airway and ependymal cilia. Furthermore, sperm from *MIS/MIS* animals showed a defect manifest as additional central apparatus microtubule not observed in airway or brain ciliary axoneme. These findings suggest that assembly of the axonemal dynein motors require different pre-assembly machinery that could not be served by the haplo-insufficient *MIS/MIS* coded protein, and that the sperm ODA motor assembly is more dependent on a fully functional DNAAF5 than other motile cilia.

Ultrastructural analysis of *Dnaaf5* mutants also informed us of activities of dynein assembly factors and of motor protein transport to the axoneme. Dnaaf5 is expressed only in the cytoplasm, and in concert with other cytoplasmic factors (e.g., *DNAAF2, SPAG1, DNAAF4, LRRC6*) participate in the assembly of cilia motor proteins. Observation in *Xenopus* suggest that IDA and ODA are assembled in unique pools (15, 55), and we saw a tendency of loss of IDAs over ODAs, indicating that Dnaaf5 is perhaps more essential for IDA assembly. Analysis of the *MIS/MIS* cilia ultrastructure also led to several unexpected observations. First, although the exact composition of the “truncated” outer arm motor structures in DNAAF5 mutant cilia are unknown, they provide sufficient cilia motility only in the *MIS/MIS* and not *MIS/NULL* genotypes. We cannot assess the longitudinal numbers of ODA to determine if the abundance is related to the MIS allele dosage. Second, in the *MIS/MIS* cilia, the presence of normal appearing outer dynein arms alongside absent or truncated outer dynein arms within the same cilia cross section is not microtubule pair specific (i.e., occurring on the same microtubule numbers 1-9), suggesting that the dynein assembly and transport machinery is both indiscriminate and may transport a partially assembled complex. Finally, assembly and transport of the dynein arm complexes may be disengaged from the microtubule assembly process (56) such that in normal conditions, both processes occur in parallel, however when one assembly factor stalls (i.e., Dnaaf5), the others continue to function, resulting in patchy assembly across the axoneme.

The observed ultrastructural changes in *Dnaaf5* mutant cilia were supported by proteomics of cilia from *MIS/NULL* mice, showing decreases in components of outer and inner dynein motor complex proteins, as well as docking proteins CCDC151 and ARMC4. Docking proteins are not known to be part of the cytoplasmic motor assembly process though may attach to the outer dynein arms during transport into the cilia (15). Other diminished proteins are those with lesser-known function, including FAP126 (57), an inner junction associated protein, and NME9, a thioredoxin-like protein predicted to be associated with motile cilia, which we previously showed interacts with DNAI2 during assembly and is present in the ciliary axoneme (55). The latter was missing in human cilia isolated from a patient with a *DNAAF5* variant, supporting the proteomic findings in *Dnaaf5* mutant cila. These findings suggest a more complex role for DNAAF5 during ciliary axoneme assembly than previously understood.

There are several limitations of our findings. First, the gene dosage effect identified may be unique to *Dnaaf5*, and additional studies are required to identify similar patterns of genotype-phenotype relationship with other cilia-associated genes. Second, the function of DNAAF5 is unknown and how the mutant proteins may interact in the axonemal motor pathway is not defined, however, we had previously shown that DNAAF5 variants continue to interact with SPAG1, a potential pathway binding partner (14). Third, while we also sought to determine if variants in *Dnaaf5* result in transcriptional responses that may provide a molecular phenotype, the use of scRNA sequencing is limited and is exploratory, with no similar reported studies for comparison. In that regard, we do not know how *MIS/MIS* mutant or the PCD subjects with *DNAAF5* variants can compensate for decreased motor proteins, though one possibility is by augmenting the overall activity within the dynein axonemal particles that are proposed to control motor protein assembly (58).

In conclusion, our data support the contention that different combinations of pathologic variants of the PCD gene *Dnaaf5* may result in varying partial function, resulting in mild or severe disease, dependent on the protein function and independent of genetic background or environment. Moreover, each allelic variant of DNAAF5 and any of the over 50 genes known to cause PCD, may lead to a gene dosage effect that contributes to the phenotype. In cases of variants that do not result in RNA decay and null conditions, interpreting the effects of compound heterozygosity of variants may require a comprehensive evaluation including functional assays to establish the contribution of each variant to disease. Such considerations do not negate the importance of environmental exposures, access to medical care and appropriate management of chronic conditions when trying to make broad genotype-phenotype links.

## Methods

### (Additional methods are in the Supplemental Information)

#### Human studies

Human studies were performed with permission from the Institutional Review Board of Washington University in Saint Louis (IRB# 201705095). Patients were seen at the Saint Louis Children’s Hospital PCD and Rare Lung Disease Clinic. Consent from parent or guardians and assent was obtained from children if old enough. Airway epithelial cells were obtained by brush of the inferior surface of the middle turbinate.

#### Generation of *Dnaaf5* mutant mice

All animal studies were performed with the permission of the Institution Animal Care and Use Committee of Washington University in Saint Louis. The *Dnaaf5* C498F mouse was created in a C57BL/6 background using reagents designed and validated at the Genome Engineering & Stem Center at Washington University. Briefly, gRNAs were designed to cleave as close to the C498 position as possible. The gRNAs were produced as synthetic CRISPR RNAs (crRNAs) that were annealed with the *trans*-acting CRISPR RNA (tracrRNA) and complexed with recombinant Cas9 protein and validated as described in **Supplemental Material**. The gRNA/Cas9 complex with both ssODNs were electroporated into single cell C57BL/6 strain embryos. Embryos (20-25) were transferred to each pseudo-pregnant C57BL/6 mother (59). Live born mice were genotyped using NGS, as during validation. Mice were contained in a microorganism barrier facility and all lines were bred in a single room. Tail biopsies were used to extract DNA, submitted to NGS, to identify genotypes.

#### Airway epithelial cell culture

Human nasal epithelial cells were isolated from biopsy brushes as previously described (14). Mouse airway epithelial cells were isolated from trachea harvested from animals and grown as previously described (36, 60). Human and mouse basal epithelial cells were expanded in culture, then differentiated on supported membranes (Transwell, Corning, NY) using air-liquid interface conditions. Cell preparations were maintained in culture for four to ten weeks.

#### Transmission electron microscopy

Fresh or cultured airway epithelial cells, sperm, and fragments of brain lateral ventricles were fixed in 2% paraformaldehyde/2% glutaraldehyde in 100 mM sodium cacodylate buffer and processed for transmission electron microscopy (TEM) as described in **Supplemental Material**. Cilia cross sections were scored by 2-3 readers, blinded to the genotype. Axonemal doublets were not scored unless the A and B microtubules could be discerned in the images.

#### Epithelial cell immunofluorescent staining

Airway cells were fixed and immunostained as previously described (36, 61). Primary antibodies used were acetylated α−tubulin (1:500, clone 6-11-B1, Sigma-Aldrich, St. Louis, MO), rabbit polyclonal DNALI1 (1:100, Sigma-Aldrich,), CFAP126 (HPA045904, 1:100, Sigma-Aldrich), and NME9 (HPA040000, 1:100 Sigma-Aldrich). Primary antibodies were detected using fluorescently labeled secondary antibodies (Alexa Fluor, Life Technologies). Nuclei were stained using 4’, 6-diamidino-2-phenylindole (DAPI). Images were acquired using a Ti2 Nikon epifluorescent microscope interfaced to a CMOS camera and Elements imaging software (Nikon, Melville, NY). Images were globally adjusted for brightness and contrast using Affinity Photo (Serif Ltd, West Bridgeford, UK).

#### High speed video-microscopy of multicilated cells

Cells were imaged live and recorded using a Nikon Eclipse Ti-U inverted microscope modified with lenses that use phase contrast and Hoffman modulation contrast (NAMC, Nikon). The microscope is enclosed in a customized environmental chamber maintained at 37 °C as described (10, 62). Images were captured by a high-speed video CMOS camera and processed with the Sisson-Ammons Video Analysis system (Ammons Engineering, Mount Morris, MI). Cilia beat frequency was analyzed in at least 5 fields obtained from each preparation, after visually confirming ciliated cells in the analyzed areas.

#### Computer-assisted sperm analysis (CASA)

Sperm obtain from the cauda epididyma were assessed assess sperm motility by CASA analysis performed using a Hamilton– Thorne digital image analyzer (HTR-CEROS II v.1.7; Hamilton–Thorne Research, Beverly, MA). Sperm capacitation tests were performed using 1 mM of 8-bromo-cAMP, with additional details in the **Supplemental Material**.

#### Tissue Histology

Mouse lungs were inflated via the trachea at 20 cm H_2_O with formalin and submerged in buffer formalin overnight at room temperature. Heads from mice were immersion fixed in Bouin’s fixative until the bones were decalcified, after which coronal slices were obtained to capture the lateral ventricles of the brain and the maxillary sinuses. Sections were stained with hematoxylin and eosin and images acquired by brightfield microscopy using a Nikon Ti2 microscope (Nikon).

#### Brain MRI imaging and ventricle volume quantification

Magnetic resonance imaging (MRI) was performed using a 4.7T Varian and 9.4T (Bruker) MRI scanner (Varian Inc., Palo Alto, CA) using settings described in the **Supplemental Materials**. The lateral and fourth ventricle volumes were calculated using ITK-SNAP software (Version 3.8.0), based on the slice thickness multiplied by the ventricle volume on each slice.

#### Cilia transport measurement

Transport on the surface of trachea and brain ventricle was determined by imaging fluorescent bead movement. Trachea were removed, fully opened across the length of the sagittal plane and submerged in a well containing PBS within a 37 °C temperature-controlled enclosure, the ciliated surface was visually confirmed. Beads (Fluoresbrite, 2 µm diameter, Polysciences, Warrington, PA) were diluted 1:500 in phosphate buffered saline, then 10µl were added to the surface of the distal trachea. For functional imaging of the ependyma cilia flow network, whole mount of the lateral wall of lateral ventricle was prepared as previously described (63). For brain studies, the diluted microbeads were deposited with a micropipette on the dorsal side, towards the anterior region of the lateral wall of the lateral ventricle. Details of flow recording are in **Supplemental Materials**.

#### Cilia Isolation and mass spectroscopy

Ciliary axonemes were isolated from the surface of highly ciliated airway cells by application of cilia buffer as described, with some modifications (64) as described in **Supplemental Material**. Tandem mass tag labeling (TMT) was performed using the TMT 10-plex reagent kit (ThermoFisher, Waltham, MA). Detailed methods of liquid chromatography-mass spectroscopy analysis is included in the **Supplemental Material**.

#### Single cell RNA sequence analysis of airway epithelial cells

Cultured primary airway cells were prepared for scRNAseq by dissociating the ALI cultures as previously described (65). Cell viability was maintained above 80% across all samples. Library preparation and sequencing was performed by the Genome Technology Access Center (GTAC) core at Washington University in St. Louis. For each sample, 20,000 cells were loaded on a Chromium Controller (10x Genomics, Pleasanton, CA) for single cell capture and cDNA was prepared according to the 10x Genomics protocols as described in **Supplemental Materials**. Paired-end sequencing reads were processed by Cell Ranger (10x Genomics software, version 2.0.0). For each sample, 5010-9910 cells were captured. The average sample had a mean of 85,446 reads per cell (ranging from 62,408 – 112,0111 depending on the library). The median number of genes detected per cell on the average sample was 4487. The samples were pre-processed using Seurat package (66), and filtered to remove stressed or dead cells (those with mitochondrial gene content of more than >25%), potential doublets (cells with more than 7500 genes detected), or low quality cells (those with less than 2500 genes detected). clustering was performed using the *FindClusters* function in Seurat. Clusters of cells were manually annotated according to their known marker gene, and gene expression and clustering results were displayed on UMAP.

#### Statistical analysis

Analysis was performed using GraphPad Prism (Version 9). Differences between two groups were compared using the Mann-Whitney U test. Multiple medians were compared using the Kruskal-Wallis test followed by Dunn’s Multiple Comparison. Paired comparisons were analyzed using the Wilcoxon Signed Rank test. The Chi-square test was used to test genotype ratios for Mendelian fit. For airway clearance studies, unpaired t tests and one-way ANOVA (Tukey’s multiple comparisons test) was performed to compare each condition. P <0.05 indicated a statistically significant difference.

## Authors contributions

A.H and S.B. designed the research studies, supervised the experiments, analyzed the data, and wrote the manuscript. A.H, D.K.G., J.X., H.X., H.X., J.H., T.H., L.D.P., S.R., S.K.B., R.M.H., S.P.G performed experiments. D.K.G contributed to RNA-seq data analysis. S.T. performed initial mass-spectrometry analysis. C.M.S. advised and supervised sperm related studies. J.M.S. advised and supervised brain related studies and provided reagents. P.M. contributed data on mouse knock phenotypes and reviewed the manuscript. M.R.M and S.K.D contributed reagents, advised on experiments and the reviewed manuscript.

## Acknowledgements

Special thanks to Heymut Omran, Munster University, Germany for advice and sharing patient-related data.

