## Supplemental material for "The effect of *Dnaaf5* gene dosage on primary ciliary dyskinesia phenotypes"

*Co-corresponding authors:

**Supplement contains:**

Supplemental Methods

Supplemental Table 1

Supplemental Figures 1-7.

### Supplemental Methods

**Generation of *Dnaaf5* mutant mice.** All animal studies were performed with the permission of the Institution Animal Care and Use Committee of Washington University in Saint Louis. The *Dnaaf5* C498F mouse was created in a C57BL/6 background using reagents designed and validated at the Genome Engineering & Stem Center at Washington University. Briefly, gRNAs were designed to cleave as close to the C498 position as possible. The gRNAs were produced as synthetic CRISPR RNAs (crRNAs) that were annealed with the *trans*-acting CRISPR RNA (tracrRNA) and complexed with recombinant Cas9 protein. The complexes were transfected into N2A cells for Next Generation Sequencing (NGS)-based validation for cleavage at the target sites. Two single stranded oligo DNA donors (ssODNs) were designed based on the more active gRNA, target sequence 5'-CACACAGCAGCAGATGCTCCAGG. ssODN1 containing the C498F mutation, as well as a silent mutation that abolishes the protospacer adjacent motif (PAM) site for the gRNA, and 60 bases of homology arms on each side was used. A second ssODN2 that only has the silent mutation that abolishes the PAM site for the gRNA, termed the block only ssODN, was also used. The gRNA/Cas9 protein complex was transfected with the ssODNs into N2A cells and validated by next generation sequencing (NGS) for efficient incorporation of the mutations before the reagents were used in for mouse zygote injection. The gRNA/Cas9 complex with both ssODNs were electroporated into single cell C57BL/6 strain embryos. Embryos (20-25) were transferred to each pseudo-pregnant C57BL/6 mother (1). Live born mice were genotyped using NGS, as during validation. Mice were contained in a microorganism barrier facility and all lines were bred in a single room. DNA was extracted from tail biopsies and submitted to NGS to identify genotypes.

**Transmission electron microscopy.** Fresh or cultured airway epithelial cells, sperm, and fragments of brain lateral ventricle were fixed in 2% paraformaldehyde/2% glutaraldehyde in 100 mM sodium cacodylate buffer for 1 hour at room temperature, then overnight at 4 °C. Samples were washed in cacodylate buffer, post-fixed in 1% osmium tetroxide (Ted Pella Inc., Redding, CA) for 1 hour at room temperature, washed in distilled water, then en block stained in 1% aqueous uranyl acetate Electron Microscopy Sciences, Hatfield, PA) for 1 hour, rinsed in distilled water, then dehydrated in a graded series of ethanol and embedded in Eponate 12 resin (Ted Pella). Sections of 95 nm were stained with uranyl acetate and lead citrate. Sections on grids were viewed on a JEOL 1200 EX transmission electron microscope (JEOL USA, Peabody, MA), images captured with an AMT 8-megapixel digital camera and AMT V602 software (Advanced Microscopy Techniques, Woburn, MA). Cilia cross sections were scored by 2-3 readers, blinded to the genotype. Axonemal doublets were not scored unless the A and B microtubules could be discerned in the images.

**High speed video-microscopy of multicilated cells.** Cells were imaged live and recorded using a Nikon inverted microscope modified with lenses that use phase contrast and Hoffman modulation contrast (NAMC; Nikon). The microscope is enclosed in a customized environmental chamber maintained at 37 °C as described (2, 3). Images were captured by a high-speed video CMOS camera and processed with the Sisson-Ammons Video Analysis system (Ammons Engineering, Mount Morris, MI). Cilia beat frequency was analyzed in at least 5 fields obtained from each preparation, after visually confirming ciliated cells in the analyzed areas.

**Computer-assisted sperm analysis (CASA).** Mice were euthanized and sperm obtain from the cauda epididyma. To assess sperm motility, 3 µL aliquots of sperm suspension was placed into a 20 microns Leja standard count 4 chamber slide, pre-warmed at 37 °C. CASA analysis was performed using a Hamilton–Thorne digital image analyzer (HTR-CEROS II v.1.7; Hamilton–Thorne Research, Beverly, MA) following analysis guidelines for CASA systems provided by the manufacturer. Phase alignment was checked and the settings used for the analysis were selected as follows: objective 1:Zeiss 10XNH; min total count: 200; frames acquired, 45; frame rate, 60 Hz; camera exposure: 8 ms; camera gain: 300; integrated time: 500 ms; elongation max%: 55; elongation min%: 1; head brightness min 55; head size max: 200 $\mu m^2$; head size min: 25 $\mu m^2$; static tail filter: false; tail brightness min: 255; tail brightness auto offset: 10; tail brightness mode: manual; progressive STR (%): 80; progressive VAP (µm/s): 50.

**Mouse sperm swim out, viability and capacitation tests.** Cauda epididymal mouse sperm were placed in 1 ml of Toyoda–Yokoyama–Hosi (TYH) medium containing: 119.30 mM NaCl, 4.70 mM KCl, 1.71 mM $CaCl_2$, 1.20 mM $KH_2PO_4$, 1.20 mM $MgSO_4$, 0.51 mM sodium pyruvate, 5.56 mM glucose, 20 mM HEPES (pH 7.4 with NaOH). After 10 min of incubation at 37 °C (swim-out), sperm suspension was collected. Concentration and initial motility after swim-out was evaluated. Viability was evaluated by Eosin-Y staining 0.5% w/v in saline. Stained spermatozoa were considered not alive. For capacitation studies, sperm recovered after swim-out were added to a buffer containing 15 mM $NaHCO_3$ and 5 mg/ml BSA. Sperm concentration was adjusted to a final maximum concentration of 10 x $10^6$ cells/mL and incubated for another 90 minutes at 37 °C for capacitation. After motility measurements, 1 mM of 8-bromo-cAMP was added and following 2 min incubation, motility was measured again.

**Airway clearance**. The clinical *Pseudomonas aeruginosa* strain PAM57-15 was prepared for airway delivery in mice (provided by T. Ferkol, Washington University). Approximately $1x10^5$ colony-forming units (CFU) was suspended in a total volume of 20 mL of sterile PBS. After oral intubation, two 10 ml volumes of bacteria were delivered into the trachea, 60 seconds apart. After 1 hour, mice were euthanized by cervical dislocation. The lungs and trachea were removed separately, homogenized, and cultured in serial dilutions in triplicates on agar plates. Following incubation at 37 °C for 16 hours, colonies were counted to determine cfu per sample. The burden of bacteria recovered in the tissue sample was determined as the mean CFU recovered, corrected for animal weight.

**Brain MRI imaging and ventricle volume quantification.** Mice were anesthetized during MRI imaging using isoflurane (2.5% induction, 1.5% maintenance). Magnetic resonance imaging (MRI) was performed using a 4.7T Varian (Varian Inc., Palo Alto, CA) and 9.4T (Bruker, Billerica, MA) MRI scanner. T2-weighted fast spin echo sequences were used to acquire images (repetition time 3000/echo time 27.50 mS, 3 averages, field of view was 18.0 mm x 18.0 mm, matrix 128 x 128, 24 axial slices, and 0.50 mm thick) with 4.7T MRI scanner and (repetition time 2852.572/echo time 66 mS, 4 averages, field of view was 14.0 mm x 14.0 mm, matrix 256 x 256, 14 axial slices, and 0.70 mm thick) with a 9.4T Bruker MRI scanner. The lateral and fourth ventricle volumes were calculated using ITK-SNAP software (Version 3.8.0), based on the slice thickness multiplied by the ventricle volume on each slice.

**Cilia transport measurement.** Transport on the surface of trachea and brain ventricle was determined by imaging fluorescent bead movement. Trachea were removed, fully opened across the length of the sagittal plane and submerged in a well containing PBS within a 37 °C temperature-controlled enclosure. The ciliated surface was visually verified by direct observation using an inverted Ti microscope. Beads (Fluoresbrite, 2 μm diameter, Polysciences, Warrington, PA) were diluted 1:500 in phosphate buffered saline, then added to the surface of the distal trachea. For functional imaging of the ependyma cilia flow network, whole mount of the lateral wall of lateral ventricle was prepared as previously described (4). For brain studies, the diluted microbeads were deposited with a micropipette on the dorsal side, towards the anterior region of the lateral wall of the lateral ventricle.

For both tissues, the flow of microbeads was recorded using a 10x objective lens and10x eyepiece using the no "zoom" function of a Nikon Ti inverted microscope. Image capture was performed at 10 frames per second (fps) for 5 seconds per image, totaling 50 frames per image. Following acquisition, files were imported to ImageJ software (Wayne Rasband, NIH, v 1.52p) and were analyzed using Trackmate (v 5.2.0) (5, 6). Image scale for calibration was set to 0.714 micrometers (pixel width and height), voxel depth 0.00 mm and time interval 0.1 sec. Particle capture was performed using the Laplacian of Gaussian (LoG) detector for detection of individual particles. This algorithm was used for estimation of the size of the particle. Parameters for "blob diameter" and "threshold" were adjusted until capture of at least 80% of beads by visual examination. Most often, a "blob diameter" of 7 mm and "threshold" of 1 were sufficient to capture most beads in the study. Another algorithm "Linear motion LAP tracker" was used to track the particles/beads with an initial search radius of 15 mm, search radius of 10 mm and 2 as maximum frame gap.

Tracks generated with beads/particles were filtered using the mean quality filter with threshold set to auto. Tracks remaining after this filtration step were used in final statistical analysis and image creation. After detection and analysis, the data was exported using "Track Statistics". The images containing color-coded tracings of the tracks were screen captured and stitched together to create a composite whole ventricle tracking image.

**Cilia Isolation for mass spectroscopy.** Ciliary axonemes were isolated from the surface of highly ciliated airway cells by application of cilia buffer as described, with some modifications (7). Cells cultured in 6-12 inserts of 12-well size Transwell, were washed with PBS then incubated for 1 min with 100 mL/well of deciliation buffer (20 mM Tris, 30 mM NaCl, 10 mM $CaCl_2$, 1 mM EDTA, pH 7.5, 0.1% TritonX100, 1 mM dithiothreitol, 0.05%, 2-mercaptoethanol) for 1 min on a vigorously moving rocker with orientation of the plate changed every 15 secs at RT. The supernatant from the wells is collected and rocking is repeated. All remaining steps are performed in the cold room and samples kept on wet ice. The supernatant from the wells was pooled, then collected at 1020g for 2 min at 4 °C to remove debris. A sample (5-10 mL) is reserved to confirm the presence of cilia later. The resulting supernatant is transferred to fresh tubes, and a cilia pellet is achieved after collection at 12,200 g for 5 min at 4 °C. The cilia pellets collect in a single tube using 200-800 mL of resuspension buffer (30 mM HEPES, 25 mM $NaCl_2$, 5 mM $MgSO_4$, 1 mM EGTA, 0.1 mM EDTA, 0.1 mM DTT, pH 7.3) to move the solution into a single. The cilia are pelleted again at 12,200 g for 5 min, at 4 °C, then repeated. The sample is finally collected at 20,000 g for 1 min at 4 °C. The remaining supernatant is carefully removed by pipet using a gel loading tip. The tube containing the pellet was snap frozen on dry ice. The presence of isolated cilia were confirmed using light microscopy and immunostaining with antibody to acetylated alpha tubulin in samples after the first isolation step to remove debris.

Just prior to mass spectroscopy analysis, the cilia pellet was suspended in 150 µL of 100 mM HEPES, pH 8.0, mixed briefly, and probe sonicated for 5 pulses. The sample was then reduced with 10 mM Tris (2-carboxyethyl) phosphine hydrochloride (TCEP) and alkylated with 25 mM iodoacetamide prior to digestion with trypsin (1:20) at 37 °C overnight. The digested sample was acidified with 1% trifluoroacidic acid, then purified with Sep-Pak C18 SPE column (Waters, Milford, MA). The extracted peptides were dried down and were then submitted for Tandem Mass Tag (TMT) labeling.

**Tandem mass tag (TMT) labeling**. Each sample was added with 50 µL of 100 mM HEPES/20% anhydrous acetonitrile, pH 8.5 buffer. Samples were labeled according to the TMT 10-plex reagent kit (ThermoFisher, Waltham, MA) per the manufacturer's instructions (only 7 channels were used in this study). Labeled digests were combined into a 2 mL microfuge tube, acidified with formic acid, subjected to Sep-Pak C18 solid phase extraction, and dried down.

**High pH reverse phase fractionation.** The dried peptide mixture was dissolved in 110 µL of mobile phase A (10 mM ammonium formate, pH 9.0) and 100 µL injected onto a 2.1 x 150 mm XSelect CSH C18 column (Waters) equilibrated with 3% mobile phase B (10 mM ammonium formate, 90% ACN). Peptides were separated using a gradient as previously described with the following parameters at a flow rate of 0.2 mL/min (8). Peptide fractions (total 96) were collected corresponding to 0.8 min each. Pooled samples were generated by concatenation in which every 12$^{th}$ fraction (i.e., 1, 13, 25, 37, 49, 61…; twelve fractions total) was combined. The pooled fractions were acidified, dried down and resuspended before injection.

**LC-MS analysis.** Each fraction was reconstituted with 25 µL of 0.1% trifluoroacetic acid. Samples (5 µl volumes) were transferred to autosampler vials for LC-MS analysis on an Orbitrap Fusion Lumos (Thermo Fisher Scientific, San Jose, CA) mass spectrometer coupled with a U3000 RSLCnano HPLC (ThermoFisher Scientific). The peptide separation was carried out on a 75 µm x 50 cm PepMap C18 column (ThermoFisher Scientific) at a flow rate of 0.3 µl/min and the following gradient: Time = 0–4 min, 2% B isocratic; 4–8 min, 2–10% B; 8–83 min, 10–25% B; 83–97 min, 25–50% B; 97–105 min, 50–98%. Mobile phase consisted of A, 0.1% formic acid; mobile phase B, 0.1% formic acid in acetonitrile. The instrument was operated in the data-dependent acquisition mode in which each MS1 scan was followed by Higher-energy collisional dissociation (HCD) of as many precursor ions in 3 second cycle (Top Speed method). The mass range for the MS1 done using Fourier Transform Mass Spectrometry (FTMS) was 375 to 1500 m/z with resolving power set to 120,000 at 400 m/z and the automatic gain control (AGC) target set to standard with a maximum fill time of 50 ms. The selected precursors were also fragmented in the FTMS using an isolation window of 0.7 m/z, a normalized AGC target of 200%, a maximum fill time of 105 ms, fixed HCD collision energy of 38%. Dynamic exclusion was performed with a repeat count of 1, exclusion duration of 30 s, and a minimum MS ion count for triggering MS/MS set to 25000 counts.

**Proteomic data analysis.** MS/MS samples were analyzed using Proteome Discoverer 2.4 (ThermoFisher Scientific). The SQUEST search engine in the Proteome Discover was set to search Mouse proteome database (Uniprot.org). The digestion enzyme was set as trypsin. The higher energy collisional dissociation (HCD) MS/MS spectra were searched with a fragment ion mass tolerance of 0.02 Da and a parent ion tolerance of 10 ppm. Oxidation of methionine and acetylation of N-terminal of protein were specified as a variable modification, while carbamidomethyl of cysteine and TMT labeling on lysine residues or peptide N-termini were specified as static modification. MS/MS based peptide and protein identifications and quantification results exported as excel sheets from Proteome Discover. Samples were further normalized and analyzed as previously described (9, 10).

**Single cell RNA sequence analysis of airway epithelial cells.** Cultured primary airway cells were prepared for scRNAseq by dissociating the ALI cultures as previously described (11). Cell viability was maintained above 80% across all samples. Library preparation and sequencing was performed by the Genome Technology Access Center (GTAC) core at Washington University in St. Louis. For each sample, 20,000 cells were loaded on a Chromium Controller (10x Genomics, Pleasanton, CA) for single cell capture and cDNA was prepared according to the 10x Genomics protocols. For sample preparation on the 10x Genomics platform, the Chromium Next GEM Single Cell 3' Kit v3.1, (PN-1000268), Chromium Next GEM Chip G Single Cell Kit, (PN-1000120), and Dual Index Kit TT Set A, (PN-1000215) were used. Purified cDNA was amplified for

11-13 cycles before being cleaned up using SPRIselect beads (Beckman-Coulter, Indianapolis, IN). cDNA concentrations were measured using an Agilent Bioanalyzer. GEX libraries were prepared as recommended by the 10x Genomics Chromium Single Cell 3' Reagent Kits User Guide (v3.1 Chemistry Dual Index) with appropriate modifications to the PCR cycles based on the calculated cDNA concentration. The concentration of each library was accurately determined through qPCR utilizing the KAPA library Quantification Kit according to the manufacturer's protocol (KAPA Biosystems, Roche, Basel, Switzerland) to produce cluster counts appropriate for the Illumina NovaSeq6000 instrument. Normalized libraries were sequenced on a NovaSeq6000 S4 Flow Cell using the XP workflow and a 50x10x16x150 sequencing recipe according to the manufacturer's protocol. A median sequencing depth of 50,000 reads/cell was targeted for each Gene Expression Library.

Paired-end sequencing reads were processed by Cell Ranger (10x Genomics software, version 2.0.0). For each sample, 5010 – 9910 cells were captured. The average sample had a mean of 85,446 reads per cell (ranging from 62,408 – 112,0111 depending on the library). The median number of genes detected per cell on the average sample was 4487. The samples were pre-processed using Seurat package (12), and filtered to remove stressed or dead cells (those with mitochondrial gene content of more than >25%), potential doublets (cells with more than 7500 genes detected), or low quality cells (those with less than 2500 genes detected). clustering was performed using the *FindClusters* function in Seurat. Clusters of cells were manually annotated according to their known marker gene, and gene expression and clustering results were displayed on UMAP.

**Supplement Table 1:** Known pathogenic variants in *DNAAF5*

| Variant | Location* | Inheritance | Reference | ClinVar ID |
|---|---|---|---|---|
| c.2394C>T (p.Arg792Ter) | HEAT 10 | Ter, homozygous | Unpublished | |
| c.2384T>C (p.Leu795Pro) | HEAT 10 | Missense, Homozygous | PMID:23040496 | 39684 |
| c.2374C>T (p.Arg792Ter) | HEAT 10 | Ter, unknown inheritance | Unpublished | 1454678 |
| c.2432-1G>C | HEAT 10 | splice mutant, Homozygous | PMID:25232951 | |
| c.2353_2352AG, (p.Ser785fs) | HEAT 10 | FS, Homozygous | PMID:29358401 | 241202 |
| c.2346C>G (p.Tyr782Ter) | HEAT 10 | Ter, unknown inheritance | Not reported | 454860 |
| c.2108_2114delinsCCACCCTGGGT (p.Met703fs) | HEAT 8 | FS, compound heterozygous | Unpublished‡ | 454858 |
| c.2083-1G>C | | homozygous | Unpublished | |
| c.2013_2028delinsATGGCC (p.Ala672fs) | HEAT 8 | FS, unknown inheritance | Unpublished | 649289 |
| c.1784-1G>T | | splice acceptor, unknown inheritance | Unpublished | 454854 |
| c.1777_1778del (p.Gln593fs) | HEAT 5-6 | FS, unknown inheritance | Unpublished | 1453546 |
| c.1731G>T (p.Trp577Cys) | HEAT 6-7 | Missense, compound heterozygous | Unpublished‡ | |
| c.1499G>T (p.C500F) | HEAT 5-6 | 1.Missense, homozygous 2.compound heterozygous | PMID:29358401 PMID:29363216† | |
| c.1142_1147delAGCTGC (p.Gln381_Leu382del) | HEAT6 | In-frame deletion, homozygous | Unpublished | |
| c.948G>A (p.Trp316Ter) | HEAT 4 | nonsense, unknown inheritance | Unpublished | 663589 |
| c.926G>A (p.Trp309Ter) | HEAT 4 | nonsense, unknown inheritance | Unpublished | 651485 |
| c.943C>T (p.Gln315Ter) | HEAT 4 | Nonsense, unknown inheritance | Unpublished | 1432731 |
| c.745del (p.His249fs) | HEAT 3 | FS, unknown inheritance | Unpublished | 1433639 |
| c.489_498del (p.His163fs) | HEAT 1-2 | FS, unknown inheritance | Unpublished | 454865 |
| c.363_373del (p.Ala122fs) | HEAT 1-2 | FS, unknown inheritance | Unpublished | 454864 |
| c.8C>T (p.Ala3Val) | N-terminus | Missense, homozygous | Unpublished | |
| c.208C>T (p.Gln70Ter) | C-terminus | nonsense, unknown inheritance | Unpublished | 843828 |
| c.55dup (p.Ala19fs) | N-terminus | FS, compound heterozygous | PMID:29363216† | 689528 |

*Location, HEAT-repeat motif site of the variant
†Reported as compound heterozygous (p.Ala1fs/Cys500Phe)
‡Reported as compound heterozygous (Met703FS/ Trp577Cys)
Abbreviations, FS, frame shift; Ter, termination

**Supplemental table 2:** Maaspectometry results of isolated cilia from MIS-NULL animals

**Supplemental table 3:** Upregulated biological processes of differentially expressed genes identified in scRNAseq of MIS-MIS primary cells

**Supplemental table 4:** Downregulated biological processes of differentially expressed genes identified in scRNAseq of MIS-MIS primary cells

### Supplemental Figures

#### Supplemental Figure 1

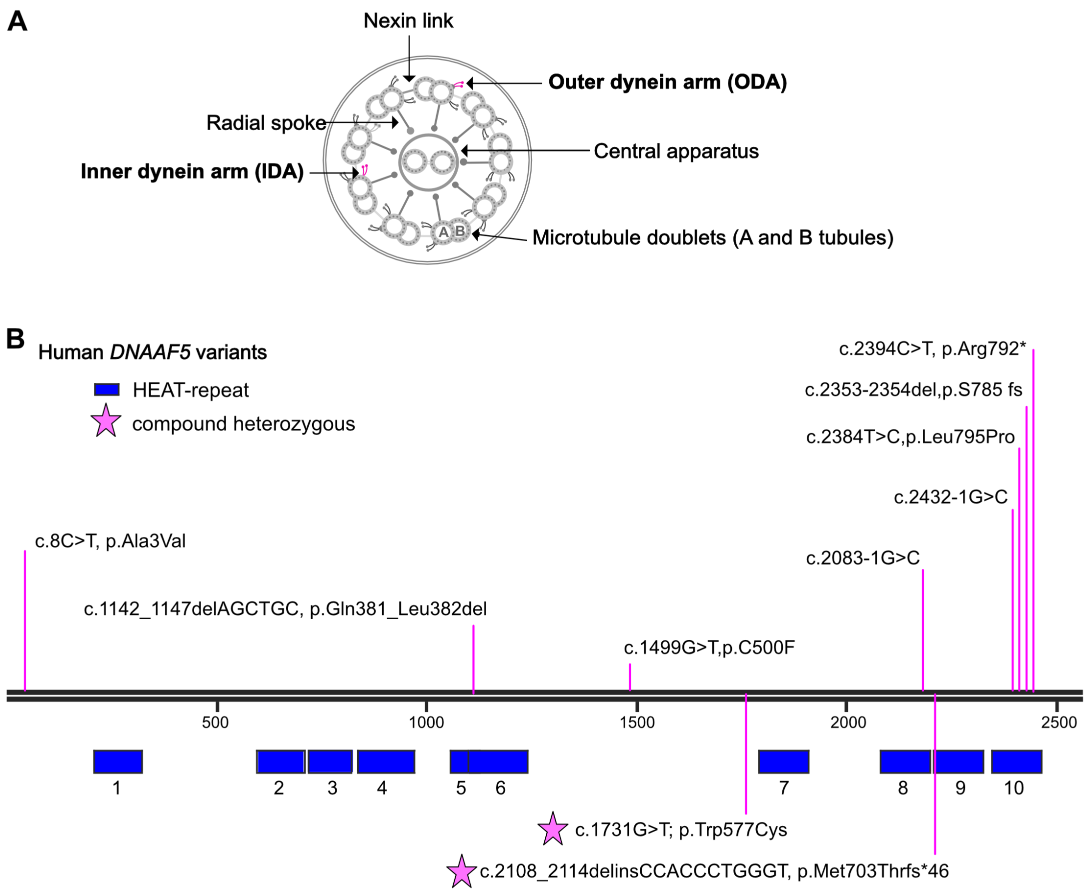


**Supplemental Figure 1. DNAAF5 variants in patients with PCD.** (**A**) Schematic of a motile ciliary cross section showing major ultrastructural features. Outer dynein arm (ODA), Inner dynein arm (IDA), Central apparatus (CA). (**B**) Identified variants in *DNAAF5* in our patient population.

**Supplement Figure 2**

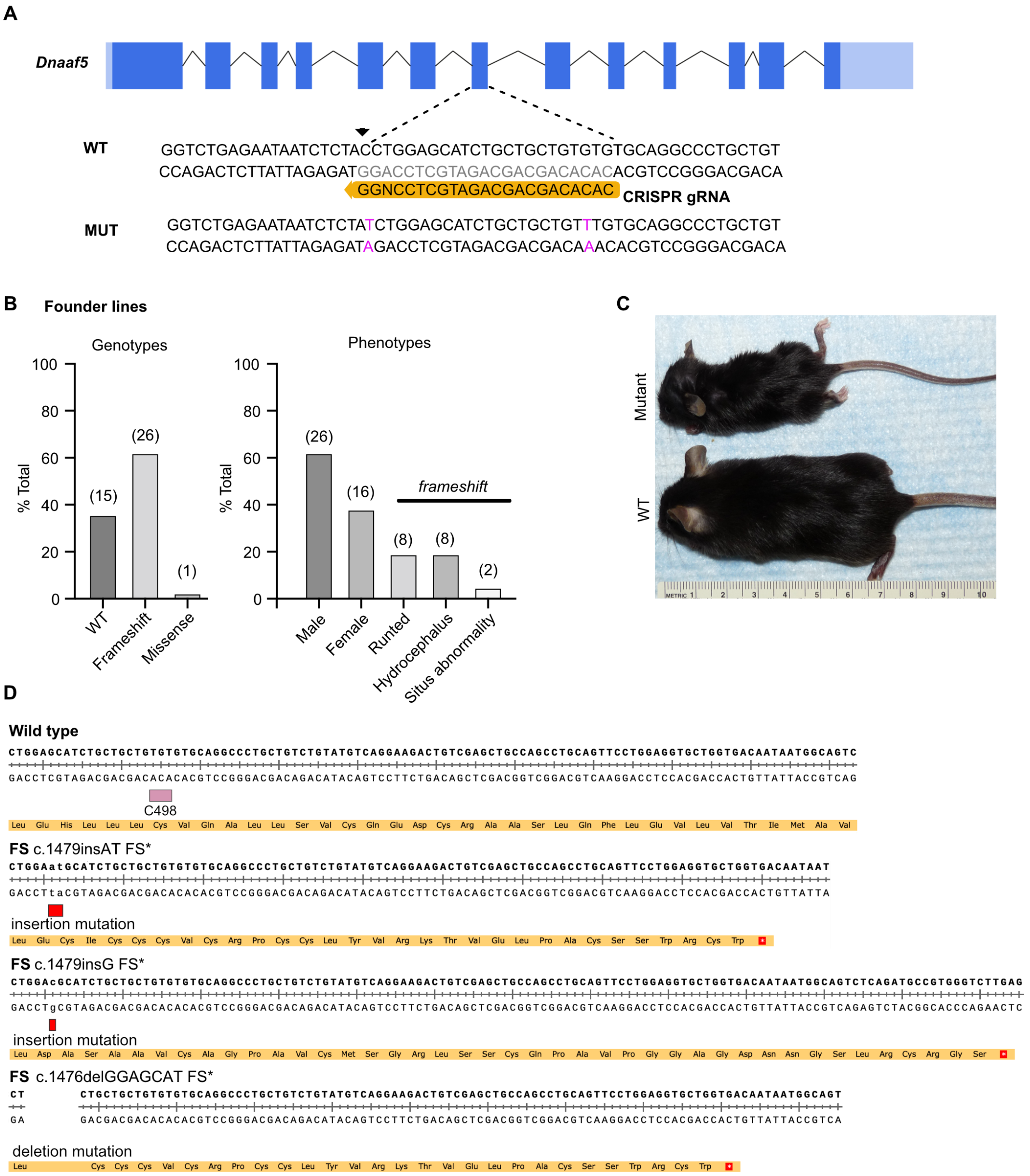


**Supplemental Figure 2. Gene editing strategy to create human ortholog *Dnaaf5* mutation.** (**A**) Gene editing strategy showing point mutation and synonymous mutation to prevent recutting by the CRIPSR-Cas9 protein. (**B**) Distribution of mutations and phenotypes identified in F0 animals. Total numbers are shown in parenthesis. (**C**) Representative example of gross morphology of F0 mutant animals, showing runted mouse. (**D**) Schematic showing frame shift mutations identified in mice after CRISPR-Cas9 editing in *Dnaaf5*.

**Supplement Figure 3**

A

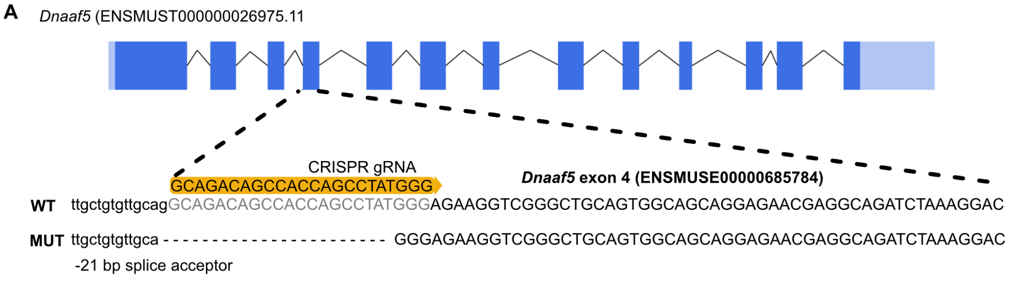


B

| NULL ALLELE | | | | |
|---|---|---|---|---|
| STAGE | WILD TYPE | HET | MUT | Litters |
| Weaning | 40 | 76 | 0 | 18 |
| E8.5 | 11 | 28 | 0 (5) | 6 |
| E6.5 | 10 | 20 | 6(2) | 5 |
| E3.5 | 11 | 17 | 9 | 4 |

Bracket denotes- partially resorbing implantation

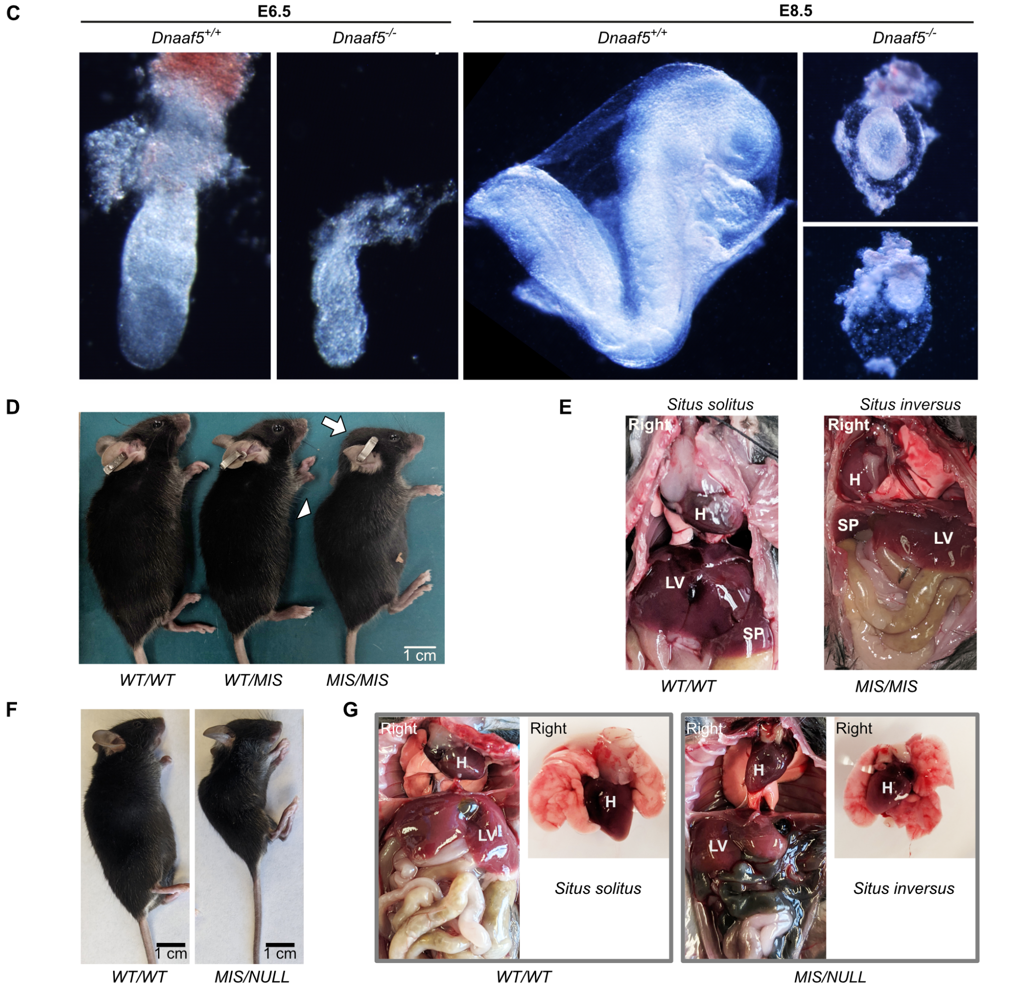


**Supplemental Figure 3. The effect of *MIS* and *NULL* alleles on gross morphology of mice. (A**) Gene editing strategy to create *Dnaaf5* knockout using CRIPSR-Cas9 protein. (**B**) Distribution on different genotypes at early embryonic day (E) showing failure of

*Dnaaf5* knockout embryos to develop. (**C**) Representative examples of Dnaaf5 knockout embryo growth arrest. (**D**) Gross morphology of *WT/WT*, *WT/MIS*, and *MIS/MIS* mice. Arrow, hydrocephalus; arrowhead, spine-deformity. (**E**) An example of *situs inversus* in *MIS/MIS* compared to *situs solitus* in *WT/WT* mice (H, heart; LV, liver; SP, spleen). (**F**) Representative images of *WT/WT* and *MIS/NULL* littermates. Note macrocephaly and lordosis in mutant animal. (**G**) Example of *situs* abnormalities in MIS/NULL animals. (H, heart; LV, liver). Side panels show the isolated heart-lung block from the same animals.

**Supplement Figure 4**

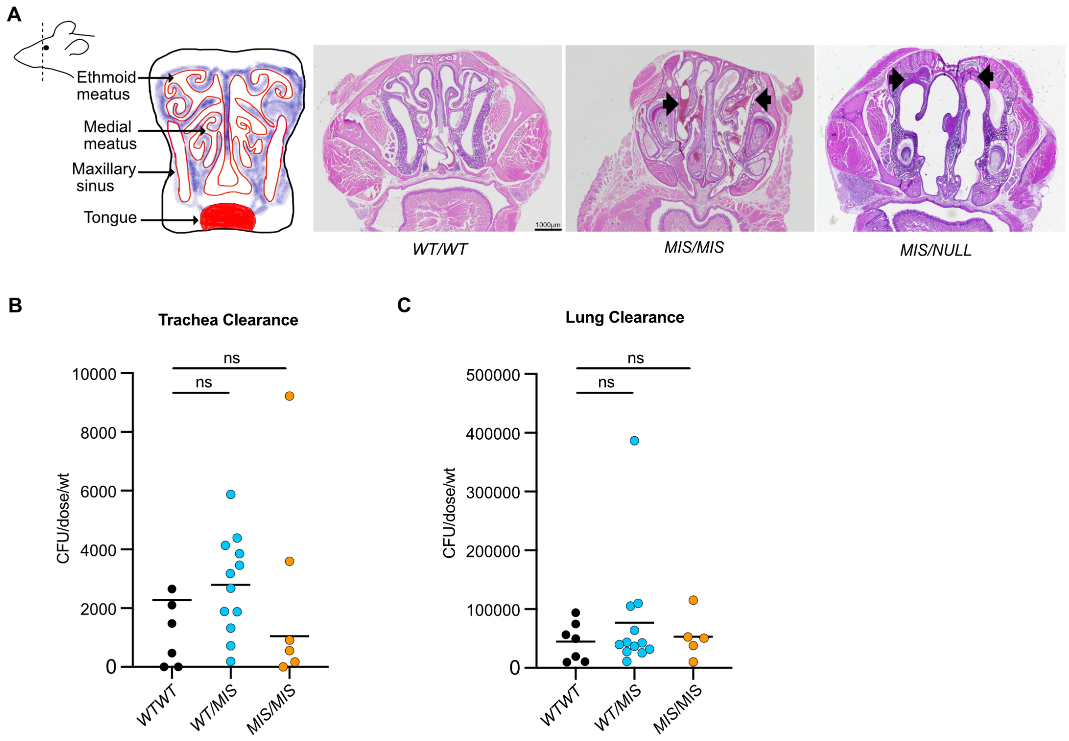


**Supplemental Figure 4. The effect of *MIS* and *NULL* alleles on sinus disease and airway clearance.** (**A**) Coronal cross sections of maxillary sinuses and nasal passages showing areas of increased inflammatory cells and mucus (arrows) in *MIS/MIS* and *MIS/NULL* animals compared to *WT/WT* littermates. Tissues were fixed and sections stained with hematoxylin and eosin (n=5-6 animals per genotype). (**B-C**) Pseudomonas clearance from trachea and lung of mutant animals showing reduced tracheal clearance trend. One hour after intratracheal delivery of approximately $10^5$ colony forming units (cfu) of Pseudomonas aeruginosa, the lungs and trachea were homogenized for bacterial culture (n=5 *WT/WT* animals, 12 *WT/MIS* animals, and 12 *MIS/MIS* animals, age 20-45 weeks). Unpaired t tests and one-way ANOVA (Tukey's multiple comparisons test) were performed to compare each condition (ns=not significant).

**Supplemental Figure 5**

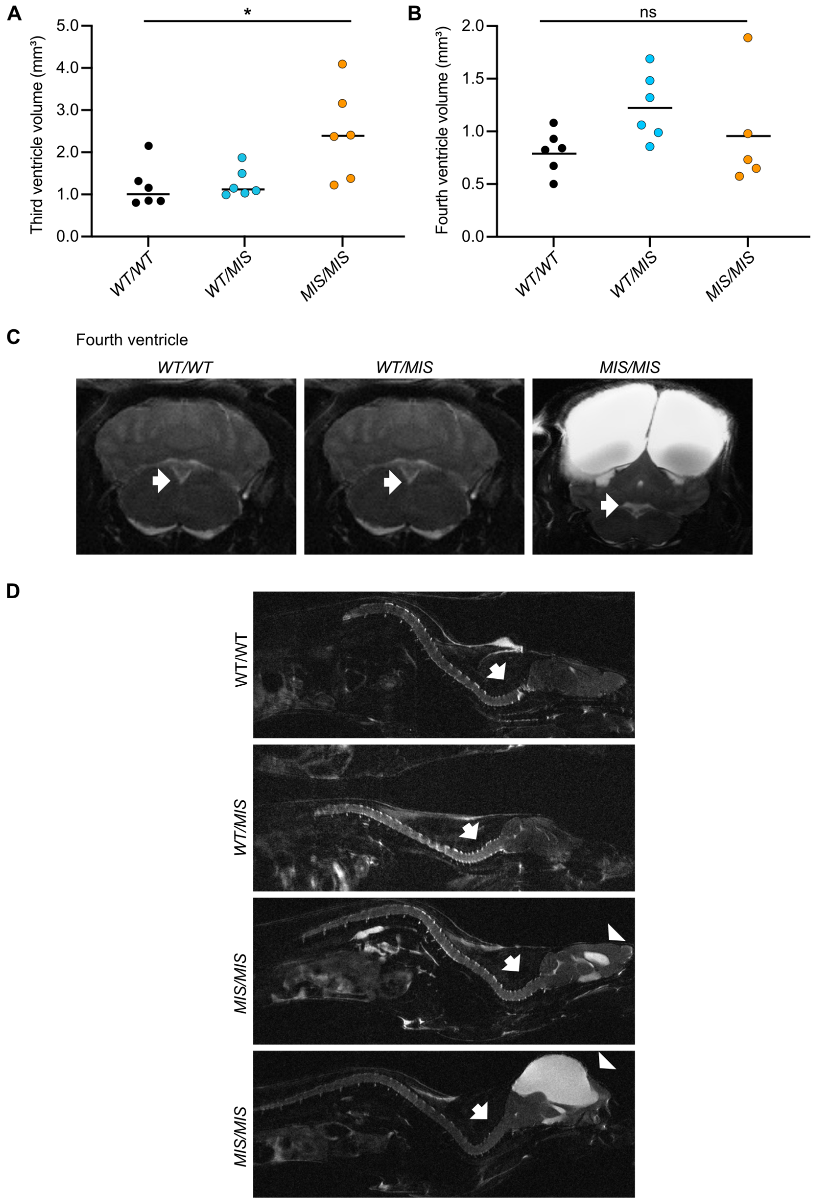


**Supplemental Figure 5 – The effect of MIS allele on brain ventricles. (A-B)** Third and fourth ventricles volume measured using MRI respectively, n=6 animals per genotype. (**C)** Representative MRI images of coronal sections brain sections showing the fourth ventricles (arrows), showing no differences in size. (**D**) Representative spine MRI images showing cervical lordosis in *MIS/MIS* animals, compared to *WT/WT* and *WT/MIS* animals. Note the cervical lordosis in *MIS/MIS* animal with minimal ventricular enlargement. Arrow indicates cervical lordosis, arrowhead indicates hydrocephalus. $^*P<0.05$, determined using Kruskal-Wallis test with Dunn's multiple comparisons; ns= not significant.

**Supplemental Figure 6**

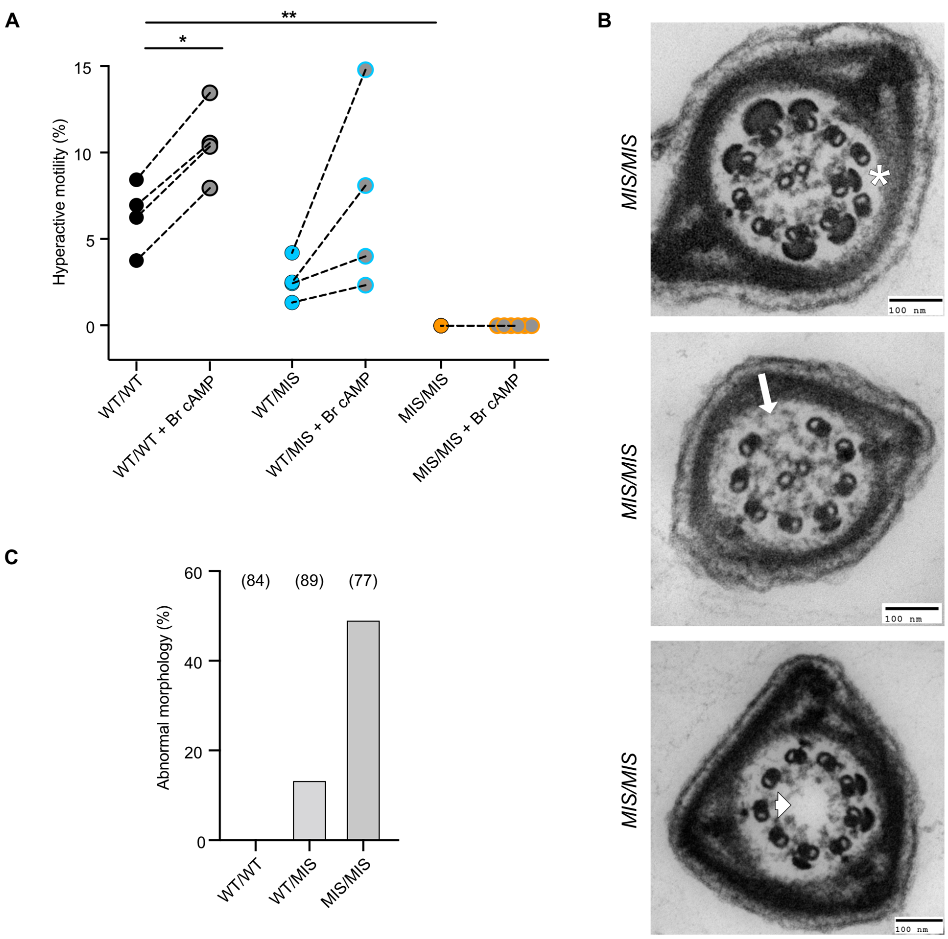


**Supplemental Figure 6. Defects in the axoneme of *Dnaaf5* mutant sperm.** (**A**) Percentage of total sperm with hyperactive (HA) motility post capacitation following induction with 8-bromo-cAMP (Br cAMP), showing an effect on *WT/WT* and *WT/MIS* sperm, but none on mutant sperm. (**B**) Examples of microtubule defects in *MIS/MIS* sperm observed on TEM. Asterisk (*) indicates extra microtubule pair, arrow indicates missing mictotubule pair, arrrowhead indicates missing central pair apparatus, (**C**) Quantification of sperm cross sections showing abnormal microtubular defects in *MIS/MIS* compared to *WT/WT* and *WT/MIS* as seen on TEM. Number of cross sections evaluated are shown in parenthesis (n=2 animals per genotype).

**Supplemental Figure 7**

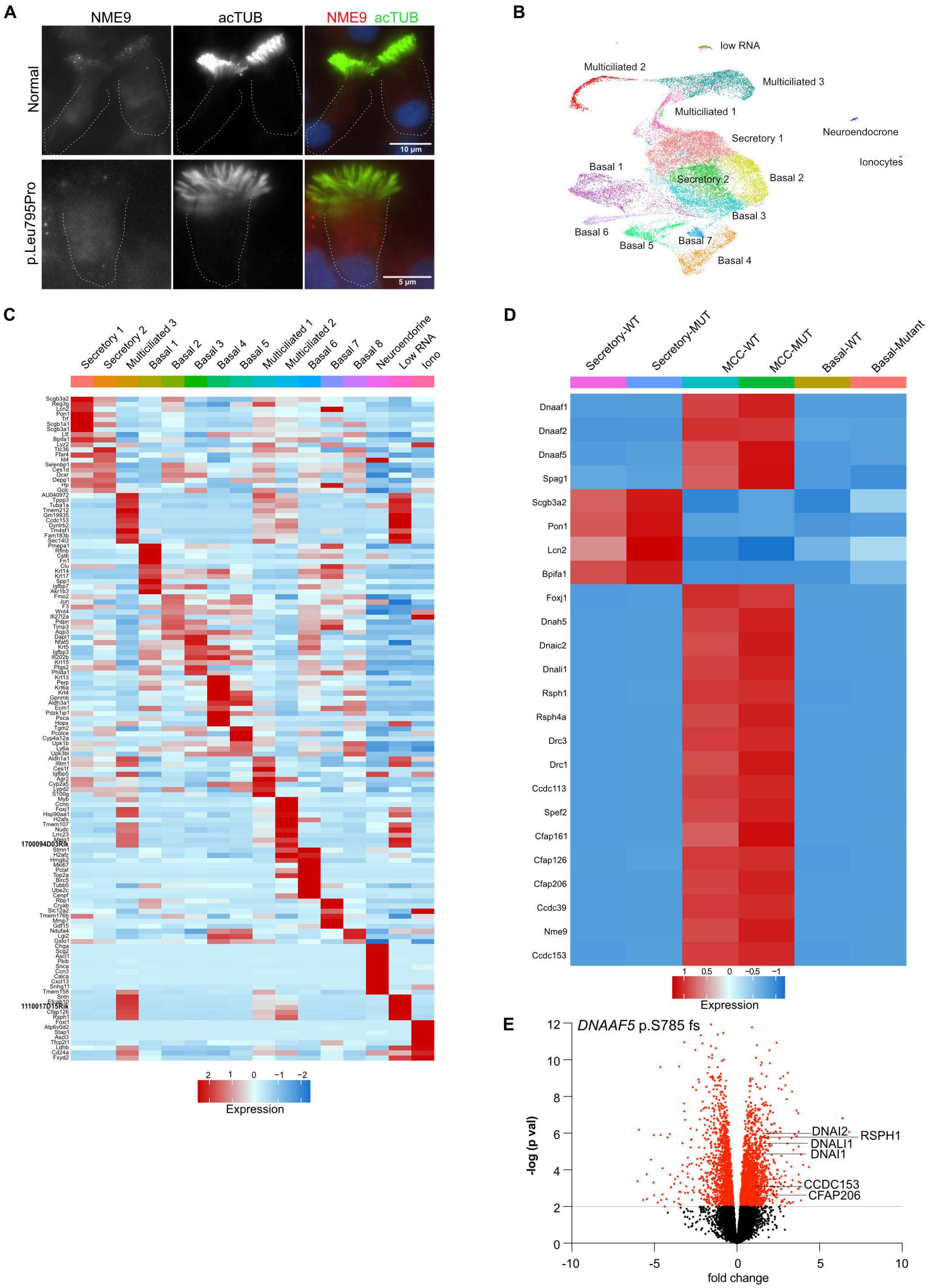


**Supplemental Figure 7. Cilia proteomics and single cell RNA sequencing analysis of *Dnaaf5* mutant mice and conserved molecular phenotypes in patients with *DNAAF5* variants.** (**A**) Immunofluorescent staining of primary culture human nasal cells showing reduced candidate protein NME9 in cilia of a patient with a *DNAAF5* variant (p. Leu795Pro). (B) UMAP showing clusters of different cell types of cultured tracheal epithelial cells from *WT/WT* and *MIS/MIS* mice. (**C**) HEAT-map showing top expressing genes in different mouse tracheal epithelial cell clusters observed in **B**. (**D**) HEAT-map showing differentially expressed cilia-related genes between *WT/WT* and *MIS/MIS* cells. (**E**) Volcano-plot of bulk RNAseq of a primary culture human nasal of a patient with a *DNAAF5* variant (p. S785fs) showing increased expression of cilia- related genes identified in *Dnaaf5* mutant mice.

**Supplemental videos**

Supplemental Video 1: Cilia beat of WT-WT primary mouse tracheal cells
Supplemental Video 2: Cilia beat of MIS-MIS primary mouse tracheal cells showing immotility.
Supplemental Video 3: Cilia beat of MIS-MIS primary mouse tracheal cells showing partial immotility.
Supplemental Video 4: Beads tracking of ex-vivo ependymal cilia of WT-WT animals.
Supplemental Video 5: Beads tracking of ex-vivo ependymal cilia of MIS-MIS animals.
Supplemental Video 6: Sperm motility of WT-WT animals.
Supplemental Video 7: Sperm motility of MIS-MIS animals.
Supplemental Video 8: Sperm tracing of WT-WT animals.
Supplemental Video 9: Sperm tracing of WT-MIS animals.
Supplemental Video10: Sperm tracing of MIS-MIS animals.
